# Exploring the Design Space of 3_10_-helical Bundles using Non-canonical Amino Acids

**DOI:** 10.64898/2026.10.16.759603

**Authors:** Xiyue Leng, Jonathan Clayden, Derek N. Woolfson

**Affiliations:** School of Chemistry, University of Bristol, Bristol, UK; Novo Nordisk Foundation Center for Protein Design, Department of Biology and Department of Drug Design and Pharmacology, University of Copenhagen, DK; Max Planck-Bristol Centre for Minimal Biology, University of Bristol, Bristol, UK; School of Biochemistry, University of Bristol, Medical Sciences Building, Bristol, UK

## Abstract

Steric constraints and favored hydrogen-bonding patterns result in peptides and proteins based on α-amino acids being largely confined to α-helical and β-strand secondary structures. Although stand-alone 3_10_ helices can be stabilized by incorporating α,α-disubstituted amino acids such as amino-isobutyric acid (Aib, `U`), assemblies based solely on this structure are rare. Previously, we reported a *de novo* designed peptide, (`ELUULK`)_4_, that forms stable and defined 3_10_-helical bundles. The Leu residues create a consolidated hydrophobic core, the Aib residues flank this core and promote 3_10_-helix formation, and the Glu/Lys residues define the water-soluble surface of the bundles. Here, we examine the mutability of these sites and their contributions to maintaining this unusual fold. Surface substitutions of Glu and Lys with Asp, Arg, or the cyclic non-canonical amino acid 4-aminopiperidine-4-carboxylic acid (Api) are well tolerated, and all variants form stable 3_10_-helical oligomers. In contrast, mutation of Aib at the flanking positions reveals strong constraints: only the cyclopropane analogue, 1-aminocyclopropane-1-carboxylic acid (Ac_3_c), preserves a 3_10_-helical bundle, albeit with reduced thermal stability. Core substitutions prove even more restrictive. Of the proteinogenic hydrophobic residues, Leu uniquely supports bundle formation. Cyclic analogues largely disrupt assembly, although 1-aminocyclopentane-1-carboxylic acid (Ac_5_c) is partially tolerated. However, extending the design to seven hexads, (`ELUULK`)_7_, enables mutation of the two central core positions to Ala or Gly and retention of stable, hexameric 3_10_-helical assemblies. These variants create hydrophobic cavities that bind environment-sensitive dyes. Together, these studies define sequence constraints for *de novo* 3_10_-helical bundles and establish routes to their functionalization.

## Introduction

*De novo* protein design has progressed from simple sequence patterning to rational and computational approaches that can specify increasingly complex structures and functions.^1–4^ Yet this progress has overwhelmingly exploited the two dominant backbone conformations of α-amino-acid polymers: α-helices and β-strands.^5, 6^ Less-common secondary structures offer different periodicities, dimensions, and modes of side-chain interactions, but their sequence-to-structure relationships remain poorly defined. The 3_10_ helix presents a stringent test of our ability to design: it occupies a region of backbone conformational space adjacent to the favored α helix, yet extended 3_10_ helices and their higher-order assemblies are rare in natural proteins.^7–11^

Experimentally observed 3_10_-helical backbone dihedral angles (φ,ψ) cluster in a restricted region near, but distinct from, the α-helical basin (Figure 1A&B). This narrow region defines the 3_10_ conformation: repetition of the φ,ψ angles generates helices with approximately three residues per turn, a 1.9 – 2.0 Å rise per residue, and a corresponding pitch of 5.8 – 6.0 Å; as a result, 3_10_ helices are stabilized by *i*→*i*+3 backbone hydrogen bonds.^8, 12, 13^ Consequently, for a given number of residues, they are longer and narrower than α helices. Thus, the 3_10_-helical geometry presents a distinct packing arrangement for forming tertiary and quaternary interactions. However, it also imposes tighter steric constraints leading to reduced stability. Consequently, even small changes in sequence and local backbone geometry can shift the balance towards the deep energy well of the neighboring α-helical state.^7, 11^

**Figure 1.**
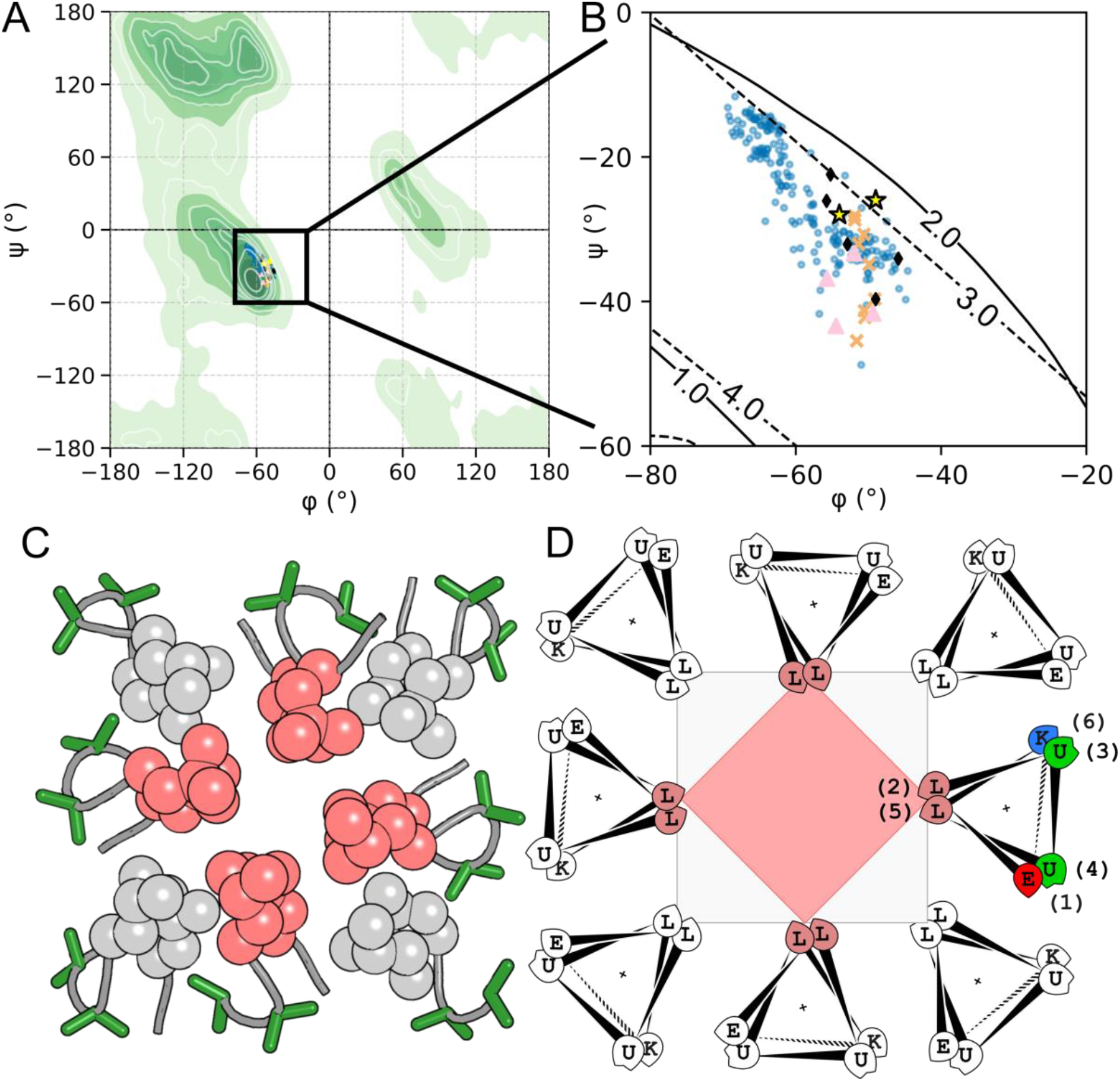
Backbone geometry and design rationale of the 3_10_-helical bundle. (**A**) Ramachandran distribution for the Top 8000 residues^19^ overlayed with backbone dihedral angles (φ, ψ) from representative 3_10_ helices. The green contour represents structural occurrence. Torsion angles were extracted using AMPAL.^20^ (**B**) Expanded region (−80° < φ < −20°; −60° < ψ < 0°) with contours of rise per residue (Å; solid) and residues per turn (dotted). The idealized 3_10_ helical geometry (3 residues per turn and 2.0 Å rise per residue) defines a narrow ridge, sampled by the experimental structures. Within the helix, all φ, ψ values are identical, raw values taken from ^21^. Key: blue dots – X-ray crystal structure of the 3_10_ peptide assembly (PDB ID:7qdi)^18^; orange crosses – CCDC entry: 780358^22^; pink triangles – CCDC entry: 982052^23^; black diamonds – CCDC entry: 982054^23^; yellow stars – canonical 3_10_ helices^17, 24^. (**C**) End-on view of the X-ray crystal structure of *de novo* 3_10_-helical bundle (PDB ID: 7qdi). The hydrophobic seam consisting of Leu side chains shown as spheres, featuring an inner (pink) and outer (gray) ring of helices. (**D**) Helical-wheel representation of the `ELUULK` hexad, identifying the solvent-exposed, flanking, and core positions (numbered 1–6). The cross marks the center of the helix.

One approach to stabilizing isolated 3_10_ helices uses α,α-disubstituted amino acids (Q4As), most notably α-aminoisobutyric acid (Aib, U),^14, 15^ Through the conformational Thorpe-Ingold effect, such residues favor compact backbone conformations.^14^ Aib-rich natural products and synthetic foldamers do form extended and stable 3_10_-helical conformations, and methods for synthesizing long, water-soluble Aib-rich peptides have made these structures accessible for design.^8, 15–17^

Previously, we have used this to design water-soluble peptides that form stable 3_10_ helix-based quaternary structure or bundles, 3_10_HD, based on a six-residue repeat (`ELUULK`)_4_.^18^ This hexad pattern matches the threefold periodicity of the 3_10_ helix: Leu residues define a hydrophobic seam, Aib flanks the seam and biases the backbone towards 3_10_ helix, and the Glu/Lys residues render a solvent-exposed charged surface. In aqueous solution, 3_10_HD is hexameric by analytical ultracentrifugation (AUC), and we solved an X-ray crystal structure for its all-D-enantiomer, which is an octameric 3_10_-helical bundle (PDB id 7qdi, Figure 1C).^18^

With this unusual 3_10_ helix-based fold established, we wanted to explore the links between sequence and structure systematically. In this paper, we report how we have probed three structurally distinct sites in the fold—the solvent-exposed charged positions, the Aib residues flanking the hydrophobic seam, and the buried hydrophobic core sites. We use proteinogenic and non-canonical amino acids (Q4As) to generate sets of designed sequences, and we combine experiments (CD, SAXS, AUC), and dipeptide torsional-energy calculations to determine and map their sequence-to-assembly relationships. Furthermore, we test if increasing peptide length can rescue destabilizing core mutations to create ligand-binding sites. Together, our results define design principles for stabilizing and tuning 3_10_-helical bundles, laying the foundation for expanding the structural repertoire of *de novo* designed peptides and protein architectures beyond α-helical space that currently dominates such designs.

## Results and discussion

### Mutability of residues in the ELUULK repeat follows a positional hierarchy

As outlined above, the `ELUULK` sequence repeat of 3_10_-HD defines three chemically and structurally distinct sites, Figure 1D: the surface-exposed Glu and Lys residues (positions 1 and 6); Aib residues flanking the hydrophobic seam (positions 3 and 4); and buried Leu residues (positions 2 and 5). We probed the mutability of these in order of increasing packing constraint, *i.e*, from the outside in.

First, for the surface sites, we replaced Glu and Lys with other charged proteinogenic amino acids, *e.g.*, aspartic acid (Asp, `D`) and arginine (Arg, `R`), or reversed their positions. Because these sites are solvent exposed, we anticipate that they would tolerate such substitutions and preserve the overall fold (Figure 1D).

The original `E–K` pairing was replaced with alternative complementary charged pairs, `D–R` and `E–R`, and by a switch in polarity, `K–D` pair, (peptides **1**–**3**, respectively; Table 1, Figure 2A). All peptides described here were synthesized by Fmoc solid-phase peptide synthesis, purified by reverse-phase HPLC, and verified by MALDI-TOF mass spectrometry (Table S1, Figures S1&S2). Biophysical characterizations were carried out in aqueous solution buffered to pH 7.4 (Table S1).

**Table 1.**
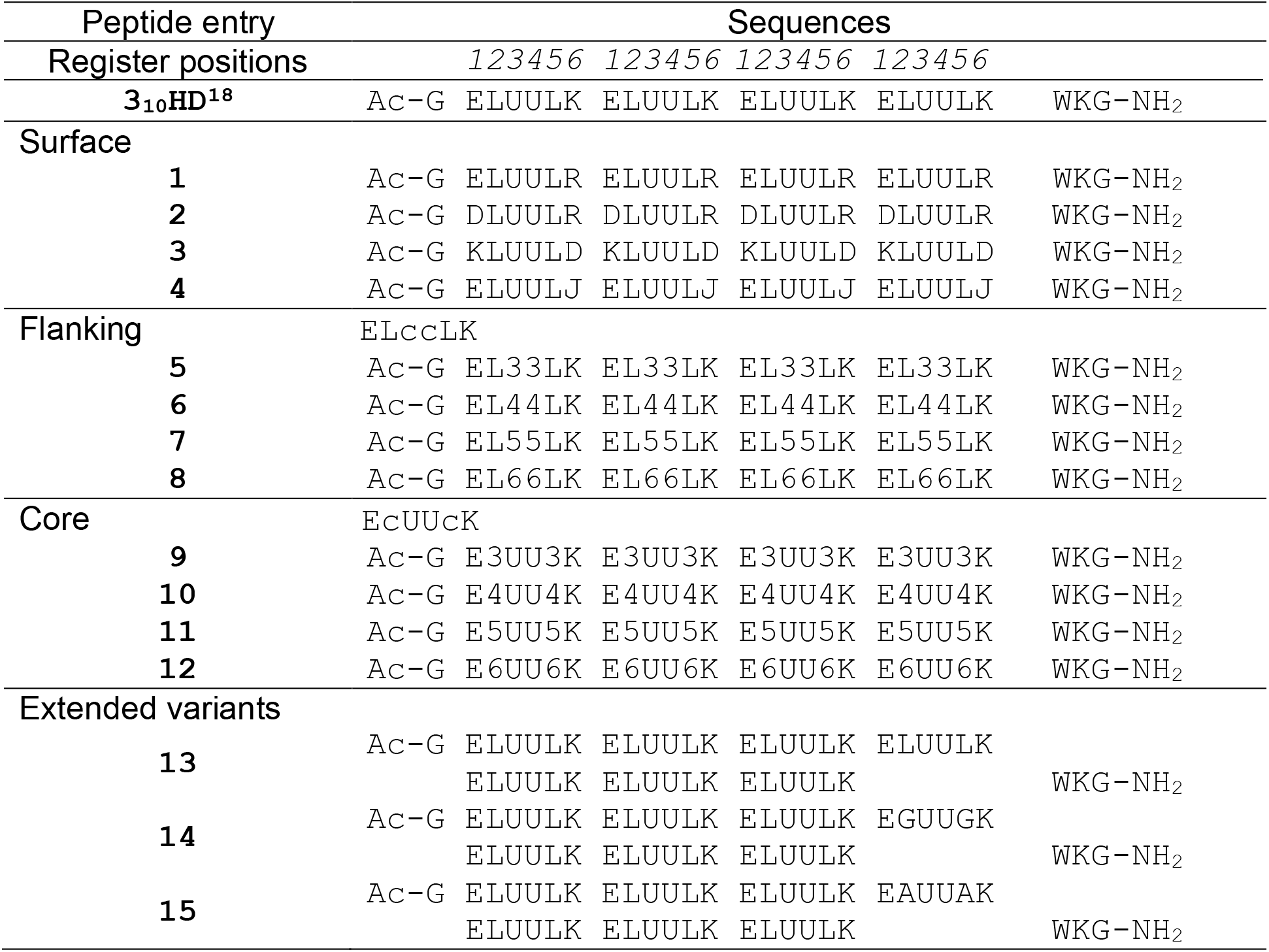
Peptide sequences discussed in this study. Non-proteinogenic residues are annotated: `U` = 2-aminoisobutyric acid (Aib), `3–6` = 3-to 6-membered 1-aminocycloalkane-1-carboxylic acids Ac_n_c (n = 3–6), `J` = 4-aminopiperidine-4-carboxylic acid (Api).

**Figure 2.**
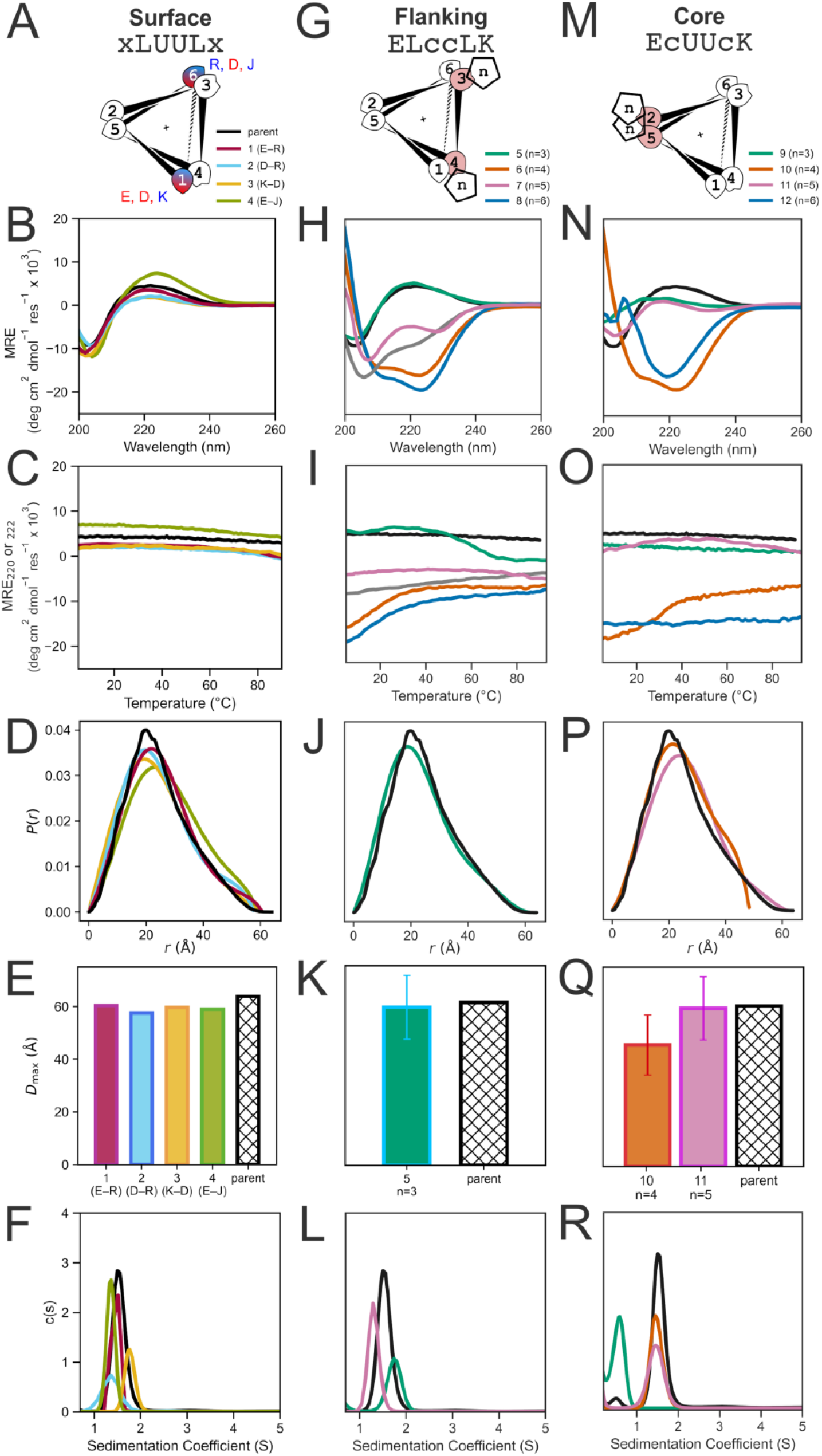
Biophysical characterization of surface (**A – F**), flanking (**G – L**), and core (**M – R**) variants, with the parent 3_10_HD peptide included as a reference throughout. (**A, G, M**) Helical-wheel representations highlighting substitutions at solvent-exposed, flanking/interfacial, and core positions, respectively. The corresponding sequence motifs are indicated above each helical wheel. (**B, H, N**) CD spectra at 5 °C. (**C, I, O**) Thermal responses of the CD signals at 220 or 222 nm (ramping from 5 – 95 °C at 1 °C min^-1^). Here, the thermal stability was assessed by monitoring 220 nm for the parent, **1 – 5**, **9**, **11**, and **12**, and 222 nm for **6 – 8**, and **10**. (**D, J, P**) SAXS pairwise distance distribution functions, P(r), obtained by indirect Fourier transformation of the Gnom-fitted SAXS data over comparable angular scattering vector q-ranges. For **1 – 4** and the parent, q_min_ = 0.017–0.022 Å^-1^, and q_max_ = 0.268–0.291 Å^-1^; for **5 – 12**, q_min_ = 0.013 Å^-1^ and q_max_ = 0.266 Å^-1^. (**E, K, Q**) Maximum particle dimensions (*D*_max_) determined from SAXS. For the surface series, *D*_max_ was 60, 58, 59, and 61 Å for **1–4**, respectively, compared with 64 Å for the parent. For the flanking series, 62 Å for **5**. For the core series, *D*_max_ was 49 Å for **10**, 63 Å for **11**, and 64 Å for the parent. The reduced *D*_max_ of **10** was consistent with its more compact α-helical assembly, whereas **11** adopted an elongated 3_10_-helical conformation with a larger rise per residue. SAXS measurements were performed at 2.5 mg ml^-1^ peptide in 20 mM Tris pH 8.0, 50 mM NaCl, 1% *v/v* glycerol. (**F, L, R**) Sedimentation-velocity analytical ultracentrifugation (AUC-SV) profiles showing the oligomeric states adopted in solution. **1 – 4** formed tetrameric, pentameric, octameric, and hexameric assemblies, respectively. Fits returned 4.3-, 4.8-, 8.0-, and 5.7-fold the monomer mass; the parent formed a hexameric assembly, corresponding to 6.2 × the monomer mass. In the flanking series, **5** and **7** formed octameric and pentameric assemblies, corresponding to 7.8 and 5.2 × the monomer mass, respectively. In the core series, **9**, **10**, and **11** formed predominantly monomeric, heptameric, and hexameric species, corresponding to 1.4-, 7.0-, and 5.9-fold the monomer mass, respectively. AUC-SV experiments were performed at 100 µM peptide in 1 × PBS, pH 7.4, at 50 krpm.

All three peptides retained the characteristic 3_10_-helical “hump–dip” far-UV circular dichroism (CD) spectrum: *i.e.*, a positive maximum at 220 nm and a negative minimum at 205 nm in a ratio of ≈1:2.^18^ These matched spectra observed for the parent design in aqueous buffer (Figure 2B and Figure S3), which is supported by an X-ray crystal structure.^16, 18^ Like the parent, none of the redesigned peptides showed a detectable thermal unfolding up to 95 °C (Figure 2C and Figure S4). Small-angle X-ray scattering (SAXS) measurements gave elongated particle-distance distributions with maximum particle dimensions (*D*_max_) of 58 – 60 Å, close to the 64 Å measured for the parent (Figure 2D&E, Table S2, and Figure S5). Analytical ultracentrifugation (AUC) revealed distinct oligomeric states: peptides **1**–**3** formed tetramer, pentamer, and octamer, respectively, compared with hexamer for the parent under the same conditions (Figure 2F, Figures S6&S7). Thus, surface-charge substitutions preserve the 3_10_-helical architecture while giving some variation in oligomeric state, analogous to the sequence-dependent oligomer tuning in α-helical coiled-coil systems.^25^

To test these positions further, we replaced the Lys residues with its cyclic quaternary analogue, 4-aminopiperidine-4-carboxylic acid (Api, `J`), to give peptide **4**, (`ELUULJ`)_4_ (Table 1, Figure 2A). This variant retained a thermally stable 3_10_-like CD spectrum and formed a discrete hexamer by AUC (Figure 2B–F, Figures S4–7). The more-intense positive band near 220 nm indicated that even a substantial backbone-constrained substitution is tolerated at these sites (Figure 2B).

Together, these data show that surface-exposed positions accommodate changes: mutations to analogous combinations of charged proteinogenic or non-canonical amino acids has little effect on 3_10_-helical assembly, while providing a handle for tuning the quaternary structure.

Second, we examined the Aib positions flanking the hydrophobic seam. Both positions 3 and 4 in all of the `ELUULK` repeats were replaced with cyclic quaternary amino acids to give (`ELccLK`)_4_, where `c` = 3-to 6-membered 1-aminocycloalkane-1-carboxylic acids, Ac_3_c–Ac_6_c (abbreviated as `3`, `4`, `5`, and `6`, peptides **5**–**8**; Table 1 and Figure 2G). This systematically varies ring size and side-chain geometry while retaining strong conformational restriction *via* the quaternary Cα centers, a feature associated with helix formation.^26–29^

Interestingly for this series, only peptide **5**, (`EL33LK`)_4_ — *i.e.*, with Aib substituted for its cyclic analogue, Ac_3_c — retained the characteristic 3_10_-helical CD spectrum and a SAXS profile consistent with an elongated assembly (*D*_max_ = 63 Å) like the parent design (Figure 2H, J&K). It unfolded reversibly with T_M_ ≈ 60 °C and formed an octamer by AUC (Figure 2I&L, Figures S6&S7). The other variants showed substantial changes in their CD profiles, consistent with predominantly α-helical species, or mixtures of these and 3_10_-helix bundles with significantly reduced T_M_ values (Figure 2H&I, Figures S3&S4). The Ac_5_c-containing peptide **7** was distinctive: its low [*θ*]_222_ / [*θ*]_208_ was consistent with literature reports of mixed or distorted 3_10_ helices^30–35^; and AUC indicated a pentamer, suggesting that steric perturbation at these flanking positions can drive alternative packing arrangements (Figures 2H&L; Figures S6&S7). By SAXS, peptides **6** and **8** did not appear to form discrete assemblies and, so, were not characterized by AUC (Figure S5).

These data indicate that the core-flanking positions tolerate some but minimal variation in side-chain geometry. For example, substitution by the cyclopropyl analogue of Aib, Ac_3_c, preserves an elongated 3_10_-helical bundle, but with reduced thermal stability and altered oligomeric state; and Ac_5_c gives a smaller assembly with only partial 3_10_-like spectral character. Thus, conformational restriction afforded by α,α-disubstituted alone is insufficient to form 3_10_-helical bundles; and the side-chain geometry must also be compatible with the compact 3₁₀-helix-helix interfaces.

Third, for the core sites, we made a final peptide series, (`EcUUcK`)_4_. Previously, we have substituted the Leu residues of (`ELUULK`)_4_ with the proteinogenic aliphatic amino acids Ala, Val, and Ile.^18^ Surprisingly, these all form partially α-helical monomers in solution. The (`EcUUcK`)_4_ series aimed to extend this and attempt to rescue 3_10_-helical structure by replacing Leu with the fully aliphatic cyclic amino acids (`3`–`6`), peptides **9**–**12**, Table 1 and Figure 2M. However, we found that these substitutions also largely disrupted the 3_10_-helical secondary structure and thermal stability (Figure 2N&O). Peptides **10** and **12** with Ac_4_c and Ac_6_c had intense α-helical and possibly β-sheet spectra, respectively (Figures S3&S4). Nonetheless, the variants either formed discrete oligomers or aggregated, suggesting that the additional α,α-disubstituted amino acids do drive towards helical assembly, but that the core packing of a 3_10_-helical bundle is frustrated (Figure 2P–R, Figures S6&S7). Among them, peptide **11** most resembled the parent, 3_10_-HD. This has the cyclopentane side chain and, therefore, the same number of carbon atoms as Leu. It retained partial 3_10_-like character CD character, formed an elongated assembly (*D*_max_ = 63 Å), and sedimented as a discrete hexamer (Figure 2N–R, Table S2 and Figures S5–7). By contrast, peptide **10**, with the smaller cyclobutane side chain, formed an α-helical heptamer in solution, Figures 2N&R; and SAXS revealed a markedly smaller *D*_max_ of 49 Å, consistent with the shorter 1.5 Å rise per residue of the α helix (Figures 2P&Q). To summarize the data from this final series of systematically redesigned peptides: of the eight aliphatic proteinogenic and α,α-disubstituted amino acids tested, only the parent Leu residue is fully tolerated in the hydrophobic core of 3_10_-HD. The substitution that gives the most similar properties to this is the cyclopentenyl side chain. Interestingly, this has the same number of carbon atoms and is analogous to Leu.

### Preferences for α- and 310-helical states can be rationalized by straightforward calculations

In an attempt to map and rationalize possible conformational changes in the redesigned peptides at the residue level, we calculated torsional free-energy surfaces for Ala, Aib, and Ac_3_c–Ac_6_c using N-acetyl and C-terminal N-methylamide-capped models (Ac–Xaa–NHMe), which allow both backbone φ and ψ angles of the central residue to be sampled (Figure 3).

**Figure 3.**
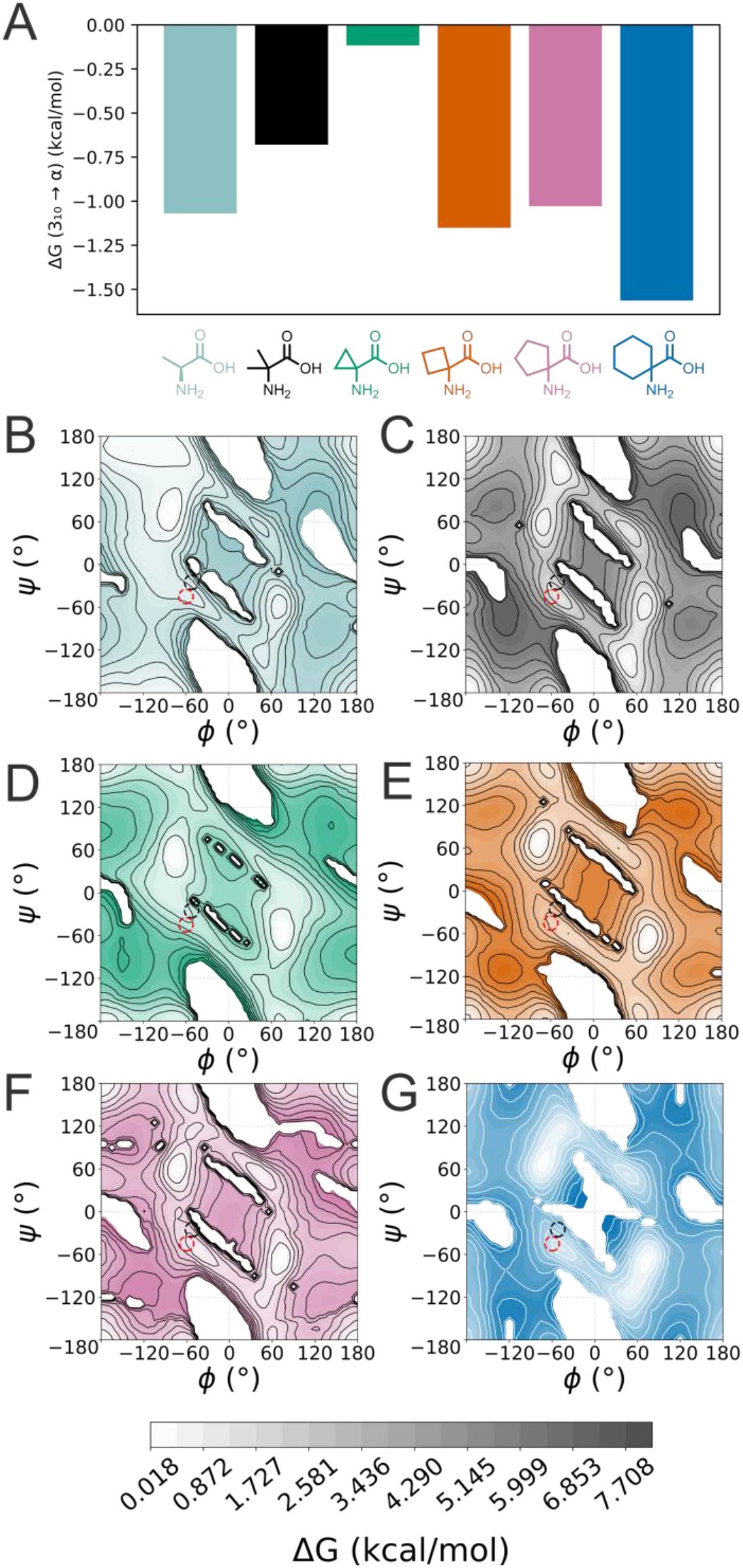
Dipeptide torsional free-energy landscapes for Ala, Aib, and cyclic Q4As. (**A**) Relative free-energy difference between the 3_10_- and α-helical basins calculated from dipeptide torsional sampling (Ac–X–NHMe) using GFN2-xTB with ALPB implicit water.^36^ The plotted sign convention is ΔG_310_◊α = G_α_ − G_310_ = −RT ln[P(α)/P(3_10_)], with conformations assigned to the 3_10_ (φ + ψ = −75 ± 10°) and α-helical (φ + ψ = −105 ± 10°) regions; values closer to zero indicate a smaller computed energetic preference for α over 3_10_. Ramachandran free-energy distributions demonstrate the backbone conformational preferences of each residue. (**B – G**) Ramachandran free-energy distributions for the dipeptides of Ala, Aib, Ac_3_c, Ac_4_c, Ac_5_c, and Ac_6_c, respectively. Ala, Aib, and the larger cyclic Q4As show pronounced low-energy minima in the α-helical region, whereas Ac_3_c gives the smallest separation between the two helical basins.

Two trends in the resulting data were striking and interesting (Figure 3A). First, as expected, Ala prefers the α-helical region, but for Aib and even more so for its cyclopropyl analogue, Ac_3_c, this preference is reduced. This is consistent with these two α,α-disubstituted amino acids partially biasing the conformational ensembles towards 3_10_-helical conformations. In turn, this will maximize the programmed amphipathic hexad sequence repeats to drive towards 3_10_-helical assemblies. Second, by contrast, all of the larger α,α-disubstituted amino acids, cyclobutyl and above, appear to return or even accentuate the preferences for the α-helical region to that of Ala (Figures 3A and B–G). These trends are broadly consistent with the experimentally observed changes in secondary structure.

### Increasing the number of hexad repeats renders mutable assemblies

The experiments above establish a gradient of tolerance in the original (`ELUULK`)_4_-based 3_10_-helical bundle design^18^, from highly permissive surface positions to apparent low mutability of core residues. Given this, we designed and tested an extended peptide with seven hexad repeats, (`ELUULK`)_7_, peptide **13** (Table 1). The aim was to use this to create a central, mutable hexad straddled by two stabilizing 3-hexad units to maintain the integrity of the folded bundle.

To test this, both Leu residues of the central hexad were changed to either glycine (Gly, G; **14**), or alanine (Ala, A; **15**), Table 1. Assuming they retain the hexameric assemblies observed for the parent, these substitutions should create cavities of ≈12 Å in length, and ≈576 – 768 Å^3^ in volume, respectively, calculated simply from the differences in side-chain volumes of Gly and Ala from Leu. Despite these significant core changes, the extended parent and both variants gave similar 3_10_-helical CD spectra (Figure 4A and Figure S3). They remained thermally stable and formed discrete hexameric assemblies by AUC (Figure 4B&C, Figures S4, S6&S7).

**Figure 4.**
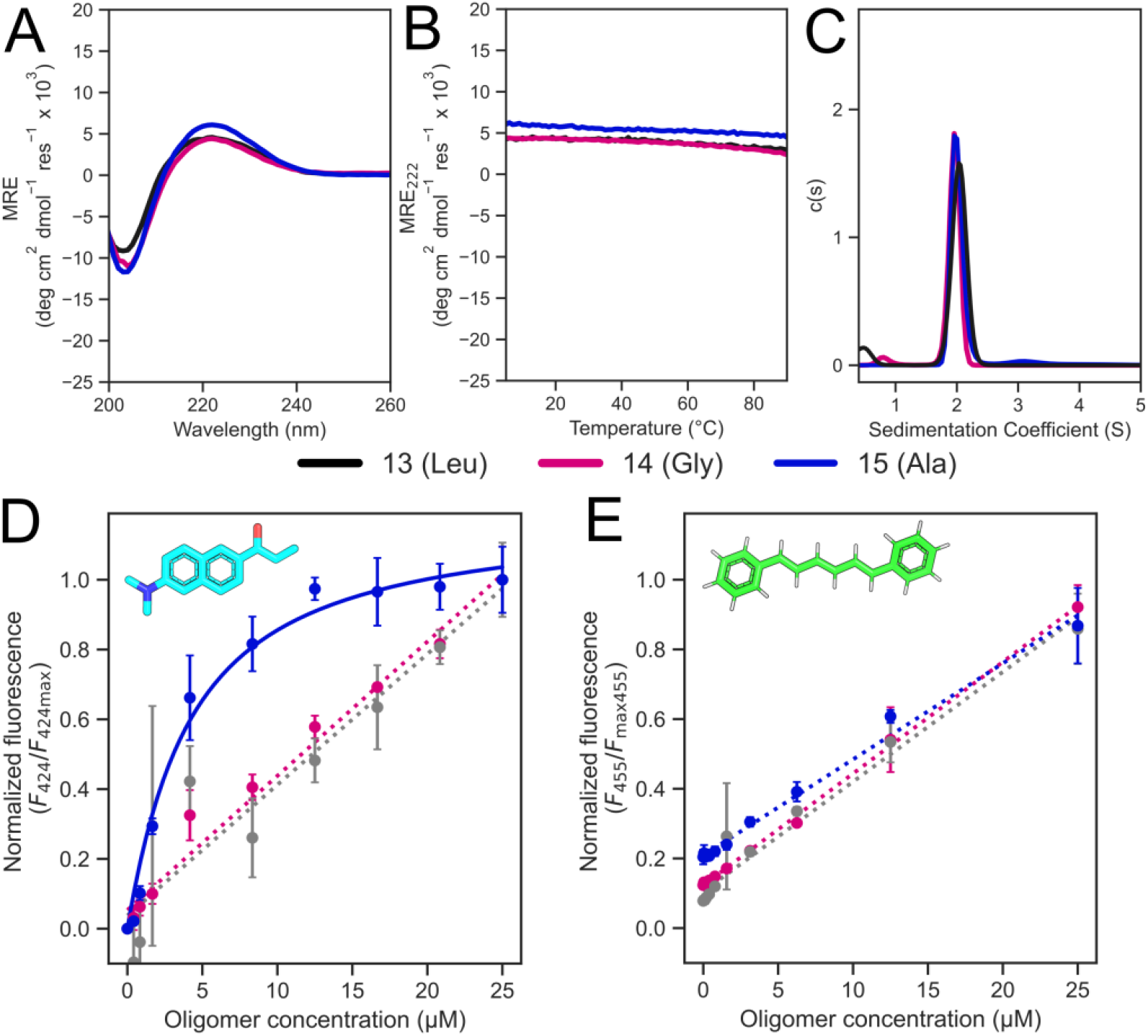
Ligand binding to extended 3_10_-helical peptide assemblies **13–15**. (**A**) CD spectra recorded at 5 °C and **(B**) thermal responses of the CD signals at 222 nm. (**C**) Overlay of AUC-SV profiles for **13** (Leu), **14** (Gly), and **15** (Ala), showing discrete hexameric assemblies in solution. Fits to the sedimentation profiles returned apparent molecular masses corresponding to 6.1, 5.6, 5.7 × the monomer masses, respectively. (**D, E**) Fluorescence-based binding assays using prodan and DPH as reporters of ligand binding. Saturation binding was observed only for **15** with prodan, giving a dissociation constant (*K*_d_) of 3.7 ± 0.8 μM (R² = 0.98). No saturation was observed for DPH binding under these conditions. Measurements were performed over 0 – 30 μM peptide oligomer with 1 μM dye in the final sample (5% *v/v* DMSO) at pH 7.4, upon incubation at 20 °C. Each dataset was normalized to its observed maximum fluorescence, and error bars represent the standard deviation of four replicate measurements. Fluorescence measurements were collected using excitation wavelengths of 371 nm for prodan and 352 nm for DPH. Saturation data were fitted with a single-site binding model.

To test for the formation of cavities in the Ala and Gly variants, we assayed for binding of the environment-sensitive dyes, 6-propionyl-2-dimethylaminonaphthalene (prodan) and 1,6-diphenylhexa-1.3,5-triene (DPH), previously used to probe for channels and cavities in *de novo* designed α-helical bundles and barrels.^37–39^ Prodan is approximately 10 × 5 Å (≈200 Å^3^), whereas DPH is ≈16 × 5 Å (≈273 Å^3^). Thus, volumetrically both should fit into the cavities, but their different lengths provide a simple test of size-and-shape-dependent interactions.

Consistent with cavity formation, both of the redesigned assemblies enhanced prodan fluorescence compared with an extended version of the parent peptide, Figure 4D. However, only the Ala variant gave a well-defined saturation curve over the concentration range tested. Anlysis returned a dissociation constant (*K*_d_) value of 3.7 ± 0.8 µM (Figure 4D, Figure S8). Notably, both variants reached comparable maximal fluorescence responses, Figure S8, suggesting that the incompletely saturated response from the Gly variant reflects weaker binding. By contrast, none of the assemblies showed saturable DPH binding under the conditions tested (Figure 4E). The stronger binding to prodan, particularly for the Ala variant, is consistent with its shorter length and better compatibility with the predicted dimensions of the cavity.

## Conclusion

In summary, the original *de novo* designed 3_10_-helical bundle, (`ELUULK`)_4_, is sensitive to mutations, but to different degrees depending on the positions and nature of the changes introduced. For example, it is robust to substitution of the surface-exposed Glu and Lys residue to other similarly charged canonical and non-canonical amino acids. Residues at these sites modulate interhelical spacing and packing, enabling some tuning of oligomeric state whilst preserving the overall 3_10_-helical architecture. By contrast, the positions flanking the hydrophobic seam impose tighter geometric constraints. From the amino acids tested, only the 2-carbon side chains of Aib and its cyclopropane analogue (Ac_3_c) are tolerated at the interfaces, indicating that subtle differences in side-chain size and conformational restriction strongly influence helix-helix packing within the compact 3_10_ geometry. The hydrophobic core positions are also very sensitive to substitution, and only tolerate Leu from the proteinogenic set of hydrophobic amino acids, or 1-aminocyclopentane-1-carboxylic acid for the cyclic set, both of which have 4 carbon atoms in their side chains.

Computational analysis of backbone conformations indicates that Aib and Ac_3_c are not simply stronger helix inducers, but are also more permissive of 3_10_ conformations: their constrained geometries reduce the energetic discrimination between the α- and 3_10_-helical backbone conformations. This provides an explanation for their abilities to retain the 3_10_-helical character in the re-designed peptides, whereas Q4As with larger cyclic side chains tend to drive towards α-helical conformations. More generally, our experimental results indicate that successful formation of the 3_10_-helical bundle requires both a compatible backbone conformations and side-chain geometries that are compatible with the packing of 3_10_ helices.

Interestingly, Ac_5_c has distinct behavior when incorporated at either buried or interfacial positions. Unlike the smaller or larger amino acids in the series, the cyclopentane ring can adopt multiple puckered conformations that relieve ring strain while maintaining compatible backbone geometries.^40^ This conformational flexibility may allow access to both 3_10_- and α-helical conformations without complete structural collapse. Therefore, its intermediate conformational preference may permit partial retention of 3_10_ helix. However, when buried in the core, its bulkier side chain appears insufficient to preserve the ideal packing arrangement of the parent Leu. This position-dependent behavior of Ac_5_c illustrates that intrinsic conformational preference alone does not determine the final structure, and the compatibility with the surrounding packing environment is important. Together, these observations suggest that the 3_10_-helical state occupies a narrow region of conformational space in which small changes in side-chain geometry can redirect its folding.

Nonetheless, despite these constraints, there is scope to mutate the hydrophobic core by extending the sequence from four to seven hexad repeats. In this way, the two Leu residues of the central hexad can be substituted for alanine or glycine to create potential hydrophobic cavities within fully folded, stable 3_10_-helical bundles. Consistent with this, these variants bind small hydrophobic dyes in the µM range. These results show that extending the scaffold can accommodate otherwise destabilizing core substitutions while preserving the global stability of the 3_10_-helical assembly. This ability to introduce defined cavities through selective removal of methylene groups, or carbon-etching, provides a strategy for building 3_10_-helical bundles as programmable scaffolds for molecular recognition. Until now, this aspect of functionalization had only been accessible for conventional α-helical and β-sheet-based protein structures, even with state-of-the-art computational methods.^41, 42^ That analogous functionality can be introduced into 3_10_-based scaffolds potentially brings them into the domain of peptide assemblies, proteins, and supramolecules as designable and functionalizable architectures.^43–45^

Thus, our findings expand the functional scope of the 3_10_ helix beyond its conventional role as a structural and transient motif. That said, the 3_10_-based scaffold is less mutation-tolerant than a canonical α-helical bundles. However, our sequence-constraint mapping, may pave the way to tune oligomerization, resolve sequence-dependent α↔3_10_ switching, and introduce a ligand-responsive internal environments. The 2 Å rise per residue of the 3_10_ helix also offers a distinct axial geometry for compact bundles and membrane spanning architectures.^46–48^ These features mean that 3_10_ helices can be used as building blocks that are complementary to α helices, rather than simply energetically less favorable analogues of these. Our resulting map may also provide benchmarks for computational methods to discriminate neighboring backbone conformations through non-canonical residue chemistry.

## Associated content

### Data availability

Experimental materials and methods, peptide sequences, purification data, biophysical and structural characterizations (PDF).

The raw data and code used in this publication have been deposited at Zenodo at DOI: 10.5281/zenodo.23316369.

## Acknowledgements

X.L. was supported by an Engineering and Physical Sciences Research Council (EPSRC)-funded PhD studentship through the Centre for Doctoral Training in Technology Enhanced Chemical Synthesis (EP/S024107/1). J.C. acknowledges funding from the ERC through Advanced Grant DOGMATRON (883786). D.N.W. is supported by the Novo Nordisk Foundation through the NNF Center for Protein Design at the University of Copenhagen (Grant number: NNF25SA0105927). We thank the Mass Spectrometry Facility for access to the EPSRC-funded Bruker UltraFlex MALDI-TOF instrument (EP/K03927X/1) at School of Chemistry, University of Bristol. We thank Diamond Light Source for access to beamline B21 (proposals mx 37593). We thank K. I. Albanese, R. Petrenas and members of the Woolfson and Clayden laboratories for many helpful discussions.

Generative AI tools were used during preparation of this manuscript to assist with language editing, including improving clarity and grammar. AI tools were not used to generate or interpret experimental data, perform scientific analysis, or determine the scientific conclusions. The authors reviewed and edited all AI-assisted text and remain fully responsible for the content, accuracy, and integrity of the manuscript.

## Notes

The authors declare no competing financial interest.

## For Table of Contents Only

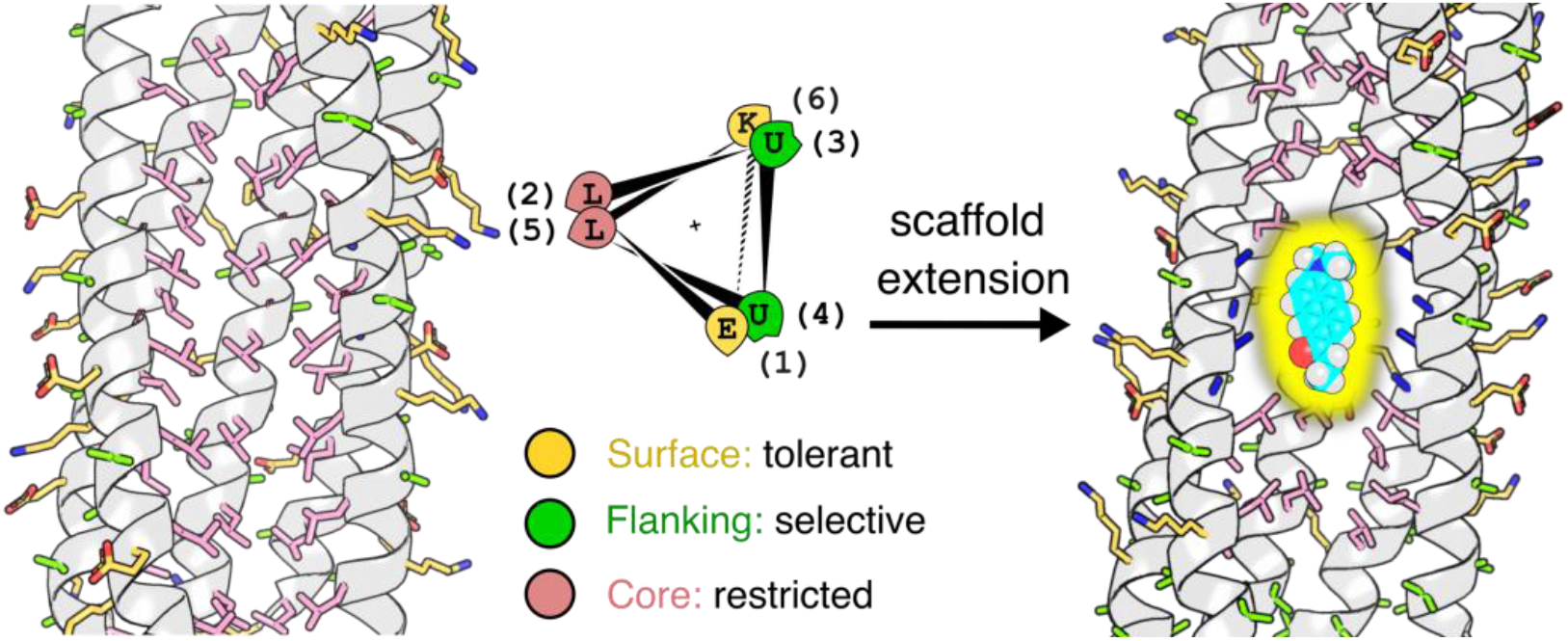

## References

(1) Albanese, K. I.; Barbe, S.; Tagami, S.; Woolfson, D. N.; Schiex, T. Computational protein design. Nat. Rev. Methods Primers 2025, 5 (13), 1–28.

(2) Pan, X.; Kortemme, T. Recent advances in de novo protein design: Principles, methods, and applications. J. Biol. Chem. 2021, 296, 100558.

(3) Woolfson, D. N. A Brief History of De Novo Protein Design: Minimal, Rational, and Computational. J. Mol. Biol. 2021, 433 (20), 167160.

(4) Kortemme, T. De novo protein design—From new structures to programmable functions. Cell 2024, 187 (3), 526–544.

(5) Woolfson, D. N. Understanding a protein fold: The physics, chemistry, and biology of α-helical coiled coils. J. Biol. Chem. 2023, 299 (4), 104579.

(6) Schweke, H.; Pacesa, M.; Levin, T.; Goverde, C. A.; Kumar, P.; Duhoo, Y.; Dornfeld, L. J.; Dubreuil, B.; Georgeon, S.; Ovchinnikov, S.;, et al. An atlas of protein homo-oligomerization across domains of life. Cell 2024, 187 (4), 999–1010.e1015.

(7) Barlow, D. J.; Thornton, J. M. Helix geometry in proteins. J. Mol. Biol. 1988, 201 (3), 601–619.

(8) Toniolo, C.; Benedetti, E. The polypeptide 310-helix. Trends Biochem. Sci. 1991, 16 (9), 350–353.

(9) Millhauser, G. L. Views of helical peptides: a proposal for the position of 3(10)-helix along the thermodynamic folding pathway. Biochemistry 1995, 34 (12), 3873–3877.

(10) Dehner, A.; Planker, E.; Gemmecker, G.; Broxterman, Q. B.; Bisson, W.; Formaggio, F.; Crisma, M.; Toniolo, C.; Kessler, H. Solution structure, dimerization, and dynamics of a lipophilic alpha/3(10)-helical, C alpha-methylated peptide. Implications for folding of membrane proteins. J. Am. Chem. Soc. 2001, 123 (27), 6678–6686.

(11) Pal, L.; Basu, G. Novel protein structural motifs containing two-turn and longer 310-helices. Protein Eng. 1999, 12 (10), 811–814.

(12) Baker, E. N.; Hubbard, R. E. Hydrogen bonding in globular proteins. Prog. Biophys. Mol. Biol. 1984, 44 (2), 97–179.

(13) Donohue, J. Hydrogen Bonded Helical Configurations of the Polypeptide Chain. Proc. Natl. Acad. Sci. 1953, 39 (6), 470–478.

(14) Toniolo, C.; Crisma, M.; Formaggio, F.; Peggion, C. Control of peptide conformation by the Thorpe-Ingold effect (Cα-tetrasubstitution). Biopolymers 2001, 60 (6), 396–419.

(15) Karle, I. L. Controls exerted by the Aib residue: helix formation and helix reversal. Biopolymers 2001, 60 (5), 351–365.

(16) Zieleniewski, F.; Woolfson, D. N.; Clayden, J. Automated solid-phase concatenation of Aib residues to form long, water-soluble, helical peptides. Chem. Commun. 2020, 56 (80), 12049–12052.

(17) Bavoso, A.; Benedetti, E.; Di Blasio, B.; Pavone, V.; Pedone, C.; Toniolo, C.; Bonora, G. M. Long polypeptide 3(10)-helices at atomic resolution. Proc. Natl. Acad. Sci. 1986, 83 (7), 1988–1992.

(18) Kumar, P.; Paterson, N. G.; Clayden, J.; Woolfson, D. N. De novo design of discrete, stable 310-helix peptide assemblies. Nature 2022, 607 (7918), 387–392.

(19) Hintze, B. J.; Lewis, S. M.; Richardson, J. S.; Richardson, D. C. MolProbity’s Ultimate Rotamer-Library Distributions for Model Validation. Proteins 2016, 84 (9), 1177–1189.

(20) Wood, C. W.; Heal, J. W.; Thomson, A. R.; Bartlett, G. J.; Ibarra, A. Á.; Brady, R. L.; Sessions, R. B.; Woolfson, D. N. ISAMBARD: an open-source computational environment for biomolecular analysis, modelling and design. Bioinformatics 2017, 33 (19), 3043–3050.

(21) Ramachandran, G. N.; Ramakrishnan, C.; Sasisekharan, V. Stereochemistry of polypeptide chain configurations. J. Mol. Biol. 1963, 7 (1), 95–99.

(22) Solà, J.; Helliwell, M.; Clayden, J. Interruption of a 3(10)-helix by a single Gly residue in a poly-Aib motif: A crystallographic study. Biopolymers 2011, 95 (1), 62–69.

(23) De Poli, M.; Byrne, L.; Brown, R. A.; Solà, J.; Castellanos, A.; Boddaert, T.; Wechsel, R.; Beadle, J. D.; Clayden, J. Engineering the Structure of an N-Terminal β-Turn To Maximize Screw-Sense Preference in Achiral Helical Peptide Chains. J. Org. Chem. 2014, 79 (10), 4659–4675.

(24) Bragg, L.; Kendrew, J. C.; Perutz, M. F. Polypeptide chain configurations in crystalline proteins. Proc. R. Soc. Lond. A Math. Phys. Sci. 1950, 203 (1074), 321– 357.

(25) Dawson, W. M.; Martin, F. J. O.; Rhys, G. G.; Shelley, K. L.; Brady, R. L.; Woolfson, D. N. Coiled coils 9-to-5: rational de novodesign of α-helical barrels with tunable oligomeric states. Chem. Sci. 2021, 12 (20), 6923–6928.

(26) Paul, P. K. C.; Sukumar, M.; Bardi, R.; Piazzesi, A. M.; Valle, G.; Toniolo, C.; Balaram, P. Stereochemically constrained peptides. Theoretical and experimental studies on the conformations of peptides containing 1-aminocyclohexanecarboxylic acid. J. Am. Chem. Soc. 1986, 108 (20), 6363–6370.

(27) Anan, K.; Demizu, Y.; Oba, M.; Kurihara, M.; Doi, M.; Suemune, H.; Tanaka, M. Helical Structures of Bicyclic α-Amino Acid Homochiral Oligomers with the Stereogenic Centers at the Side-Chain Fused-Ring Junctions. Helv. Chim. Acta 2012, 95 (10), 1694–1713.

(28) Demizu, Y.; Doi, M.; Kurihara, M.; Maruyama, T.; Suemune, H.; Tanaka, M. One-Handed Helical Screw Direction of Homopeptide Foldamer Exclusively Induced by Cyclic α-Amino Acid Side-Chain Chiral Centers. Chem. Eur. J. 2012, 18 (8), 2430– 2439.

(29) Benedetti, E.; Di Blasio, B.; Pavone, V.; Pedone, C.; Santini, A.; Crisma, M.; Valle, G.; Toniolo, C. Structural versatility of peptides from Cα,αdialkylated glycines: Linear Ac3c homo-oligopeptides. Biopolymers 1989, 28 (1), 175–184.

(30) Toniolo, C.; Formaggio, F.; Tognon, S.; Broxterman, Q. B.; Kaptein, B.; Huang, R.; Setnicka, V.; Keiderling, T. A.; McColl, I. H.; Hecht, L.;, et al. The complete chirospectroscopic signature of the peptide 310-helix in aqueous solution. Biopolymers 2004, 75 (1), 32–45.

(31) Silva, R. A.; Yasui, S. C.; Kubelka, J.; Formaggio, F.; Crisma, M.; Toniolo, C.; Keiderling, T. A. Discriminating 3(10)-from alpha-helices: vibrational and electronic CD and IR absorption study of related Aib-containing oligopeptides. Biopolymers 2002, 65 (4), 229–243.

(32) Polese, A.; Formaggio, F.; Crisma, M.; Valle, G.; Toniolo, C.; Bonora, G. M.; Broxterman, Q. B.; Kamphuis, J. Peptide Helices as Rigid Molecular Rulers: A Conformational Study of Isotactic Homopeptides from α-Methyl-α-isopropylglycine, [L-(αMe)Val]n. Chem. Eur. J. 1996, 2 (9), 1104–1111.

(33) Crisma, M.; Bisson, W.; Formaggio, F.; Broxterman, Q. B.; Toniolo, C. Factors governing 310-helix vs α-helix formation in peptides: Percentage of Cα-tetrasubstituted α-amino acid residues and sequence dependence. Biopolymers 2002, 64 (5), 236–245.

(34) Formaggio, F.; Crisma, M.; Rossi, P.; Scrimin, P.; Kaptein, B.; Broxterman, Q. B.; Kamphuis, J.; Toniolo, C. The First Water-Soluble 310-Helical Peptides. Chem. Eur. J. 2000, 6 (24), 4498–4504.

(35) Oba, M.; Nonaka, H.; Doi, M.; Tanaka, M. Conformational studies on peptides having dipropylglycine (Dpg) or 1-aminocycloheptanecarboxylic acid (Ac7c) within the sequence of l-leucine (Leu) residues. J. Pept. Sci. 2016, 106 (2), 210–218.

(36) Bannwarth, C.; Ehlert, S.; Grimme, S. GFN2-xTB─An Accurate and Broadly Parametrized Self-Consistent Tight-Binding Quantum Chemical Method with Multipole Electrostatics and Density-Dependent Dispersion Contributions. J. Chem. Theory Comput. 2019, 15 (3), 1652–1671.

(37) Parasassi, T.; Krasnowska, E.; Bagatolli, L.; Gratton, E. LAURDAN and PRODAN as polarity-sensitive fluorescent membrane probes. J. Fluoresc. 1998, 8, 365–373.

(38) Thomas, F.; Dawson, W. M.; Lang, E. J. M.; Burton, A. J.; Bartlett, G. J.; Rhys, G. G.; Mulholland, A. J.; Woolfson, D. N. De Novo-Designed α-Helical Barrels as Receptors for Small Molecules. ACS Synth. Biol. 2018, 7 (7), 1808–1816.

(39) Dawson, W. M.; Shelley, K. L.; Fletcher, J. M.; Scott, D. A.; Lombardi, L.; Rhys, G. G.; LaGambina, T. J.; Obst, U.; Burton, A. J.; Cross, J. A.;, et al. Differential sensing with arrays of de novo designed peptide assemblies. Nat. Commun. 2023, 14 (1), 383.

(40) Aleman, C.; Zanuy, D.; Casanovas, J.; Cativiela, C.; Nussinov, R. Backbone conformational preferences and pseudorotational ring puckering of 1-aminocyclopentane-1-carboxylic acid. J. Phys. Chem. B 2006, 110 (42), 21264– 21271.

(41) An, L.; Said, M.; Tran, L.; Majumder, S.; Goreshnik, I.; Lee, G. R.; Juergens, D.; Dauparas, J.; Anishchenko, I.; Coventry, B.;, et al. Binding and sensing diverse small molecules using shape-complementary pseudocycles. Science 2024, 385 (6706), 276–282.

(42) Zhu, J.; Liang, M.; Sun, K.; Wei, Y.; Guo, R.; Zhang, L.; Shi, J.; Ma, D.; Hu, Q.; Huang, G.;, et al. De novo design of transmembrane fluorescence-activating proteins. Nature 2025, 640 (8057), 249–257.

(43) Chandramouli, N.; Ferrand, Y.; Lautrette, G.; Kauffmann, B.; Mackereth, C. D.; Laguerre, M.; Dubreuil, D.; Huc, I. Iterative design of a helically folded aromatic oligoamide sequence for the selective encapsulation of fructose. Nat. Chem. 2015, 7 (4), 334–341.

(44) Collie, G. W.; Bailly, R.; Pulka-Ziach, K.; Lombardo, C. M.; Mauran, L.; Taib-Maamar, N.; Dessolin, J.; Mackereth, C. D.; Guichard, G. Molecular Recognition within the Cavity of a Foldamer Helix Bundle: Encapsulation of Primary Alcohols in Aqueous Conditions. J. Am. Chem. Soc. 2017, 139 (17), 6128–6137.

(45) Wang, P. S. P.; Nguyen, J. B.; Schepartz, A. Design and High-Resolution Structure of a β3-Peptide Bundle Catalyst. J. Am. Chem. Soc. 2014, 136 (19), 6810–6813.

(46) Jones, J. E.; Diemer, V.; Adam, C.; Raftery, J.; Ruscoe, R. E.; Sengel, J. T.; Wallace, M. I.; Bader, A.; Cockroft, S. L.; Clayden, J.;, et al. Length-Dependent Formation of Transmembrane Pores by 310-Helical α-Aminoisobutyric Acid Foldamers. J. Am. Chem. Soc. 2016, 138 (2), 688–695.

(47) Futaki, S.; Fukuda, M.; Omote, M.; Yamauchi, K.; Yagami, T.; Niwa, M.; Sugiura, Y. Alamethicin−Leucine Zipper Hybrid Peptide: A Prototype for the Design of Artificial Receptors and Ion Channels. J. Am. Chem. Soc. 2001, 123 (49), 12127–12134.

(48) Nagao, T.; Mishima, D.; Javkhlantugs, N.; Wang, J.; Ishioka, D.; Yokota, K.; Norisada, K.; Kawamura, I.; Ueda, K.; Naito, A. Structure and orientation of antibiotic peptide alamethicin in phospholipid bilayers as revealed by chemical shift oscillation analysis of solid state nuclear magnetic resonance and molecular dynamics simulation. Biochim. Biophys. Acta - Biomembr. 2015, 1848 (11A), 2789–2798.

