## supplementary information pdf for "Exploring the Design Space of 3_10_-helical Bundles using Non-canonical Amino Acids"

### Methods

#### General

All solvents, chemicals, and reagents were purchased from commercial sources and used without further purification. Fluorenylmethoxycarbonyl (Fmoc)-α-L-amino acids, Rink amide MBHA resin for solid-phase peptide synthesis (SPPS) and N,N-dimethylformamide (DMF) were purchased from Sigma-Aldrich and Cambridge Reagents. Coupling reagents Oxyma Pure and diisopropylcarbodiimide (DIC) were purchased from Fluorochem and Sigma-Aldrich, respectively. Morpholine and trifluoroacetic acid (TFA) were purchased from Sigma-Aldrich; pyridine was from Thermo Fisher; triisopropylsilane (TIPS) was from Acros Organics. All other chemicals were reagent grade and purchased from Sigma-Aldrich.

#### Peptide synthesis and purification

Automated microwave SPPS was performed on a Liberty Blue (CEM) synthesizer with inline UV monitoring. The synthesis was performed with side-chain protection of the amino acids as follows: Glu(O$^{t}$Bu), Lys(Boc), Trp(Boc). Rink amide (MBHA, 0.68 mmol/g loading, 100-200 mesh) was used for a synthesis on a 0.1 mmol scale. The coupling reactions were performed by adding amino acids dissolved in DMF (2.5 mL, 0.2 M), the coupling reagent DIC in DMF (1.0 mL, 1 M) and Oxyma Pure in DMF (1 mL, 0.5 M) to the respective resin. Standard couplings were performed at 90 °C for 4.5 min (100 W for 20 s, 60 W for 10 s, 35 W for 240 s). Standard deprotections were performed using 20% (*v/v*) morpholine in DMF at 90 °C for 1.5 min (125 W for 30 s, 32 W for 60 s). The coupling reactions concerning Q4As and their *N*-

terminal adjacent residues were carried out following the SPPS method reported by our groups.[1]

All peptides were manually acetyl capped through addition of pyridine (0.5 mL) and acetic anhydride (0.25 mL) in DMF (9.25 mL), shaking at room temperature for 20 min. The resin was washed 3 times with DMF followed by 6 times with $CH_2Cl_2$ before cleavage. Peptides were cleaved from the resin with addition of 10 mL of a mixture 95:2.5:2.5 *v/v* TFA / $H_2O$ / TIPS, shaking at room temperature for 2 h. The TFA solution was then filtered to remove the resin beads and was reduced in volume to ≈ 5 mL or lower using a flow of $N_2$. Cleaved peptide was precipitated with cold diethyl ether (≈ 45 mL), isolated *via* centrifugation and dissolved in a 1:1 mixture MeCN/$H_2O$. Crude peptides were lyophilized to yield a white or off-white powder. Crude peptides were kept in -20 °C for short-term storage, and -80 °C for long-term storage.

All peptides were purified by preparative scale high-performance liquid chromatography (HPLC, JASCO) using a Luna C18 (Phenomenex) reverse-phase column (150 × 10 $mm^2$, 5 μM particle size, 100 Å pore size). Crude peptide was dissolved at 7 mg $mL^{-1}$ in 20% *v/v* MeCN in $H_2O$ with 0.1% TFA, injected onto the column and eluted with a 3 mL $min^{-1}$ linear gradient (20-100%) of MeCN in $H_2O$ with 0.1% TFA each over 40 min. Elution of the peptide was detected with inline UV monitoring at 220 and 280 nm wavelengths simultaneously. A column oven (50 °C) was employed to improve separation. Pure fractions were identified by analytical HPLC and matrix-assisted laser desorption/ionization–time of flight (MALDI-TOF) mass spectrometry, then pooled, and lyophilized.

Analytical HPLC traces were obtained using a Jasco 2000 series HPLC system and a Phenomenex Kinetex C18 (100 × 4.6 $mm^2$, 5 μM particle size, 100 Å pore size) reverse-phase column. Elution of the peptide was detected with inline UV monitoring at 220 and 280 nm wavelengths simultaneously. The linear gradient was 20–100% *v/v* MeCN in $H_2O$ with 0.1% TFA each over 25 min at a flow rate of 1 mL $min^{-1}$. When required, a column oven (50 °C) was employed to assist peptide elution.

**Mass spectrometry**

Mass spectra were collected on a Bruker UltraFlex MALDI-TOF mass spectrometer operating in positive-ion reflector mode. Peptides were spotted on a ground steel target plate using α-cyano-4-hydroxycinnamic acid dissolved in 1:1 *v/v* MeCN/$H_2O$ with 0.1% TFA as the matrix. Masses quoted are for the monoisotopic mass as the singly protonated species.

**Solution-phase biophysical characterisation**

<u>Peptide concentration determination</u>

Peptide concentration was determined at 280 nm using a Nanodrop 2000 (ThermoScientific) spectrometer ($\varepsilon_{280}$(Trp) = 5690 $cm^{-1}$) or an Agilent (Stockport, UK) Cary 100 UV/vis spectrophotometer. For sequences without Trp, concentrations were determined using the amide absorbance for each sequence, using values determined by Gruppen and Kuipers[2] using the equation, where $\varepsilon_{peptide}$ bond is 923 $M^{-1}cm^{-1}$ , and $\varepsilon_{amino}$ acid corresponds to the molar extinction coefficient of the free amino acid at 214 nm:

$$\varepsilon_{\text{peptide}}(M^{-1}\text{cm}^{-1}) = \varepsilon_{\text{peptide bond}} \times n_{\text{peptide bonds}} + \sum_{i=1}^{20} \varepsilon_{\text{amino acid}(i)} \times n_{\text{amino acid}(i)}$$

#### Circular dichroism

Circular dichroism data were collected on a JASCO J-810 or J-815 spectropolarimeter fitted with a Peltier temperature controller in the far UV region. Spectra Manager (1.55) was used for data collection. Circular dichroism spectra were acquired as 100 μM overall assembly solution in phosphate buffered saline (PBS; 8.2 mM sodium phosphate dibasic, 1.8 mM potassium phosphate monobasic, 137 mM sodium chloride, 2.4 mM potassium chloride), pH 7.4 at 5 °C. Data were collected in a 1 mm quartz cuvette between wavelengths of 190 nm and 260 nm with the instrument set as follows: band width, 1 nm; data pitch, 1 nm; scanning speed, 100 nm $min^{-1}$; response time, 1 s. Each circular dichroism spectrum was obtained by averaging eight scans and subtracting the background signal of the buffer and cuvette. For thermal response experiments, the circular dichroism signal at wavelength of 220 or 222 nm was monitored over the temperature range 5–95 °C at a ramping rate of 1 °C $min^{-1}$ with the same settings and peptide concentrations given above.

The spectra were converted from ellipticities (mdeg) to mean residue ellipticities (MRE, $deg \cdot cm^2 \cdot dmol^{-1} \cdot res^{-1}$) using Equation 1. [3]

$$MRE = \frac{\theta \times 10^6}{c \times l \times n} \qquad \text{Equation 1}$$

where the variable $\theta$ is the measured difference in absorbed circularly-polarized light in millidegrees, $c$ is the micromolar concentration of the compound, $l$ is the path length of the cuvette in mm, and $n$ is the number of amide bonds in the polypeptide.

#### Analytical ultracentrifugation (AUC)

Analytical ultracentrifugation (AUC) was performed on a Beckman Optima X-LI or X-LA analytical ultracentrifuge with an An-50-Ti or An-60-Ti rotor (Beckman-Coulter).

Buffer densities, viscosities and peptide partial specific volumes ($\bar{v}$) were calculated using SEDNTERP (http://rasmb.org/sednterp/). However, SEDNTERP cannot recognize ncAAs. Hence, for their $\bar{v}$ calculations, pairs of Aib were replaced by Ala + Val.

The mass of the sedimenting species and oligomeric state (*i.e.*, mass observed divided by the monomer mass) of a peptide sample can be determined using sedimentation velocity (SV) and Sedimentation equilibrium (SE) experiments using the Svedberg equation (Equation 2)[4],

$$s = \frac{m(1-\bar{v}p)}{f} \qquad \text{Equation 2}$$

where s is the sedimentation coefficient in seconds (commonly expressed in Svedberg unit, S; 1 S = $10^{-13}$ s), m is the mass of the particle, $\bar{v}$ is the partial specific volume of the sample (mL/g), p is the buffer density, *f* is the frictional coefficient which depends on the shape and size of the particle (elongated or rough particles are expected to have a larger *f*, α helical coiled-coil bundles of similar lengths are expected to have a frictional coefficient around 1.2).[5] Centrifugation applies a sedimenting force to particles in solution, this is opposed by drag, buoyancy and back-diffusion. The combination of all of them is the effective centrifugal force felt by the particle. Combining the Svedberg equation with the Einstein relation (Equation 3):

$$D = \frac{RT}{f} \qquad \text{Equation 3}$$

arrives at the molecular-weight specific description of the Svedberg equation (Equation 4):

$$\frac{s}{D} = \frac{M_p(1-\bar{v}p)}{RT} \qquad \text{Equation 4}$$

where D is the diffusion coefficient ($cm^2$/s), $M_p$ is the mass of the sedimenting species $m = \frac{M_p}{N_A}$, R is the gas constant, and T is the absolute temperature (K). $M_p(1 - \bar{v}p)$

represents the effective mass of the particle (*i.e.*, the mass remaining after accounting for buoyant force, based on Archimedes' principle). The absorbance across the radial distance of the cell is monitored as a proxy for concentration distribution of the solute. The sedimentation boundary (the moving front between the cleared buffer and the sedimentation zone that marks particle migration) was monitored over time.

For SV experiments, peptide sample solutions of 310 or 410 µL were prepared in PBS at sample concentration of 250 µM and placed in a sedimentation velocity cell with an epon or aluminium, 2-channel centrepiece and sapphire windows. The reference channel was loaded with 320 µL (or 420 µL, respectively) of PBS buffer. The samples were centrifuged at 60 krpm at 20 °C, with absorbance scans taken across a radial range of 5.8–7.3 cm at 5 min intervals to a total of 120 scans. Data from a single run were fitted to a continuous $c(s)$ distribution model using SEDFIT, at 95% confidence level. Residuals for sedimentation velocity experiments are shown as a bitmap in which the grayscale shade indicates the difference between the fit and raw data. Scans are ordered vertically, with earlier scans at the top. The horizontal axis is the radial range over which the data were fitted.

SE experiments were centrifuged at speeds in the range of 20-45 krpm and scans at each recorded speed were duplicated after equilibration for 8 h. The sedimentation force and back diffusion reach a dynamic balance by allowing the sample to be centrifuged at a constant speed for an extended period of time. The resulting steady-state concentration gradient is monitored across the radial distance of the cell and modelled to determine molecular weights. SE experiments were performed in triplicate in a six-channel centerpiece (110 µL per channel) channels at 20 °C. The samples data were fitted using SEDPHAT to a single species model. Monte Carlo analysis was performed to obtain fits with 95% confidence limits.

**Small angle X-ray scattering**

SAXS data was collected at the Diamond Light Source beamline B21 using batch mode. Peptide samples were prepared at concentrations at 0.5, 1, 1.5, 2 mg $mL^{-1}$ in 20 mM Tris, 50 mM NaCl, 1% *v/v* glycerol at pH 8.0. Buffer subtraction and data merging was performed using ATSAS.[6, 7] The overall shape was further investigated by performing *ab initio* modelling using Dammif (20 models) assuming P1 symmetry.[6] Quality of fit was assessed by calculating $\chi^2$ between experimental scattering profiles to Dammif models.

**Ligand binding**

Ligand-binding assays were performed in quadruplicate using an epMotion 5070 liquid handler (Eppendorf). The total concentration of the dye was held constant at 1 µM dye in PBS with 5% *v/v* DMSO. The concentration of the *de novo* peptide assembly varied from 0–30 µM, assuming all formed hexameric assemblies. Peptide and ligand were incubated for 2 h with shaking,

Fluorescence data was collected on a Clariostar plate reader (BMG Labtech, 5.40 R3) at 20 °C, using excitation wavelengths of 371 nm for prodan, and 352 nm for DPH. The binding constants were determined by fitting the data to the following equation:

$$y = B_{max} \frac{(c + x + K_d) - \sqrt{(c + x + K_d)^2 - 4cx}}{2c}$$

Where c is the total concentration of the constant component (dye), x is the concentration of the variable component (peptide assembly), $B_{max}$ is the fluorescence signal when all of the constant component is bound, and y is the fluorescence intensity.

### Supplementary Data

#### Supplementary Tables

Table S1 Summary of biophysical characterization of peptides discussed in this study

| Peptide entry | Secondary structure | $T_M$ (°C) | SV weight (Observed / expected) | SE weight (Observed / expected) | Mass (Da) |
|---|---|---|---|---|---|
| parent | $3_{10}$ | > 95 | 6.2 | 6.3 | 3358.1 |
| **1** | $3_{10}$ | > 95 | 4.3 | 4.0 | 3051.8 |
| **2** | $3_{10}$ | > 95 | 4.8 | 4.8 | 3044.8 |
| **3** | $3_{10}$ | > 95 | 8.0 | 6.2 | 3156.9 |
| **4** | $3_{10}$ | > 95 | 5.5 | n.d. | 3212.9 |
| **5** | $3_{10}$ | 58 | 7.8 | 7.5 | 3087.9 |
| **6** | α | 25 | n.d. | n.d. | 3198.9 |
| **7** | α / $3_{10}$ | > 95 | 5.2 | 4.8 | 3309.9 |
| **8** | α | 20 | n.d. | n.d. | 3421.2 |
| **9** | α / $3_{10}$ | > 95 | 1.4 | n.d. | 2860.5 |
| **10** | α | 25 | 7.0 | n.d. | 2972.6 |
| **11** | α / $3_{10}$ | > 95 | 6.0 | 6.1 | 3084.6 |
| **12** | mixed | > 95 | n.d. | n.d. | 3196.6 |
| **13** | $3_{10}$ | > 95 | 6.1 | 6.2 | 4709.2 |
| **14** | $3_{10}$ | > 95 | 5.6 | 6.1 | 4980.1 |
| **15** | $3_{10}$ | > 95 | 5.7 | 5.8 | 4952.0 |

Table S2 SAXS statistics for peptides discussed in this study

| | 1 | 2 | 3 | 4 | 5 |
|---|---|---|---|---|---|
| **Guinier analysis** | | | | | |
| $I$(0) ($cm^{-1}$) | 0.01251 | 0.01612 | 0.00647 | 0.02854 | 0.02134 |
| $R_g$ (Å) | 19.45 | 18.99 | 18.57 | 21.03 | 18.48 |
| **P(r) analysis** | | | | | |
| $I$(0) ($cm^{-1}$) | 0.01251 | 0.01612 | 0.00647 | 0.02854 | 0.02134 |
| $R_g$ (Å) | 19.46 | 19.03 | 18.58 | 21.03 | 18.49 |
| $D_{max}$ (Å) | 61 | 58 | 60 | 59 | 62 |
| q range ($Å^{-1}$) | 0.0171 - 0.2770 | 0.0273 - 0.2908 | 0.0140 - 0.2581 | 0.0203 - 0.2868 | 0.0130 - 0.2511 |

| | 10 | 11 | 13 | 14 | 15 |
|---|---|---|---|---|---|
| **Guinier analysis** | | | | | |
| $I$(0) ($cm^{-1}$) | 0.01765 | 0.02885 | 0.03311 | 0.01178 | 0.04438 |
| $R_g$ (Å) | 18.67 | 19.91 | 27.41 | 26.97 | 27.20 |
| **P(r) analysis** | | | | | |
| $I$(0) ($cm^{-1}$) | 0.01765 | 0.02885 | 0.03311 | 0.01178 | 0.04438 |
| $R_g$ (Å) | 18.67 | 19.91 | 27.45 | 27.00 | 27.27 |
| $D_{max}$ (Å) | 48 | 63 | 93 | 91 | 94 |
| q range ($Å^{-1}$) | 0.0193 - 0.2636 | 0.0163 - 0.2495 | 0.0121 - 0.3010 | 0.0107 - 0.2501 | 0.0153 - 0.2703 |

#### Supplementary Figures

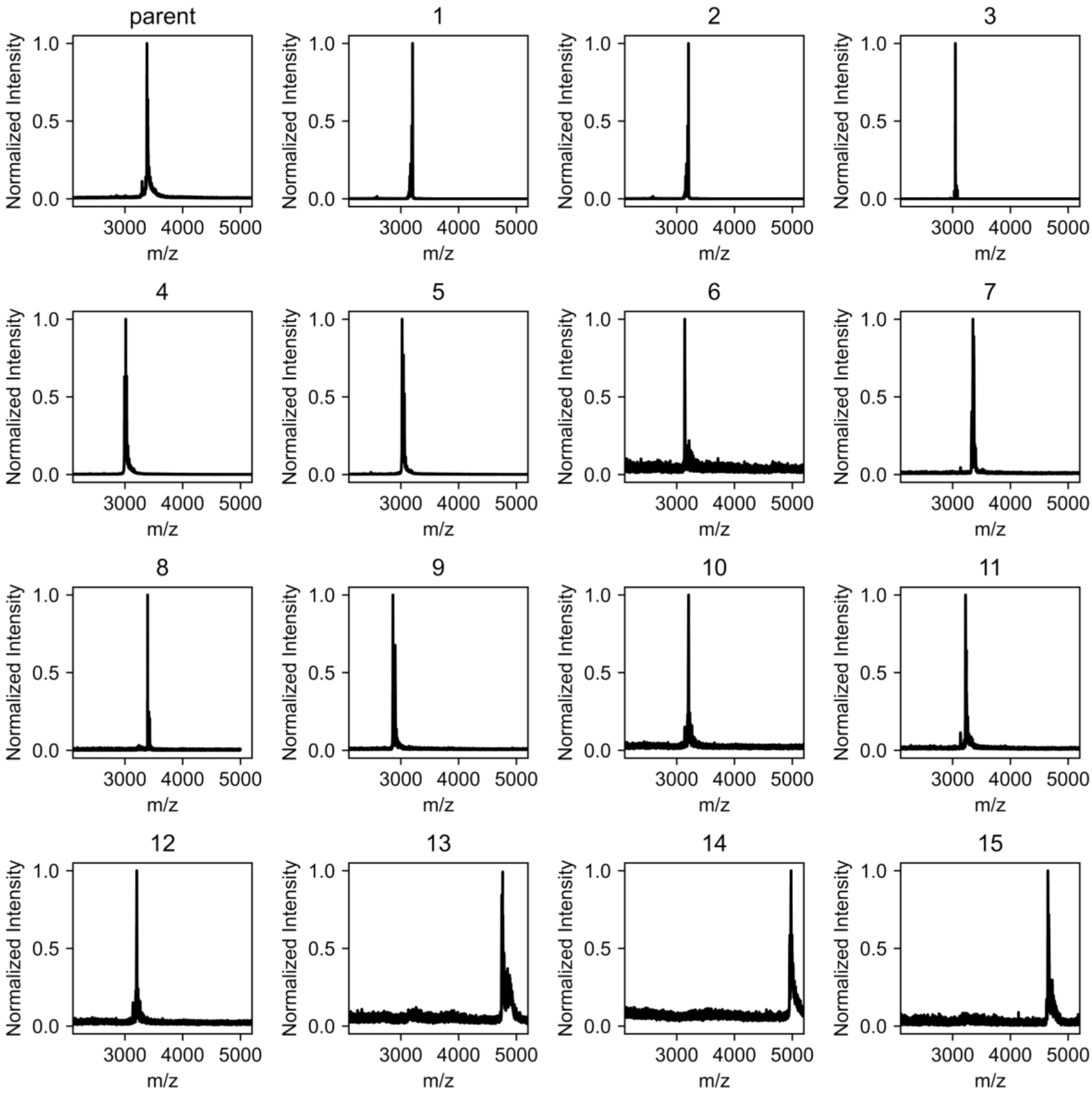


Figure S1 MALDI-TOF spectra of *de novo* peptides described in this work. Mass for individual peptides can be found in Table S1.

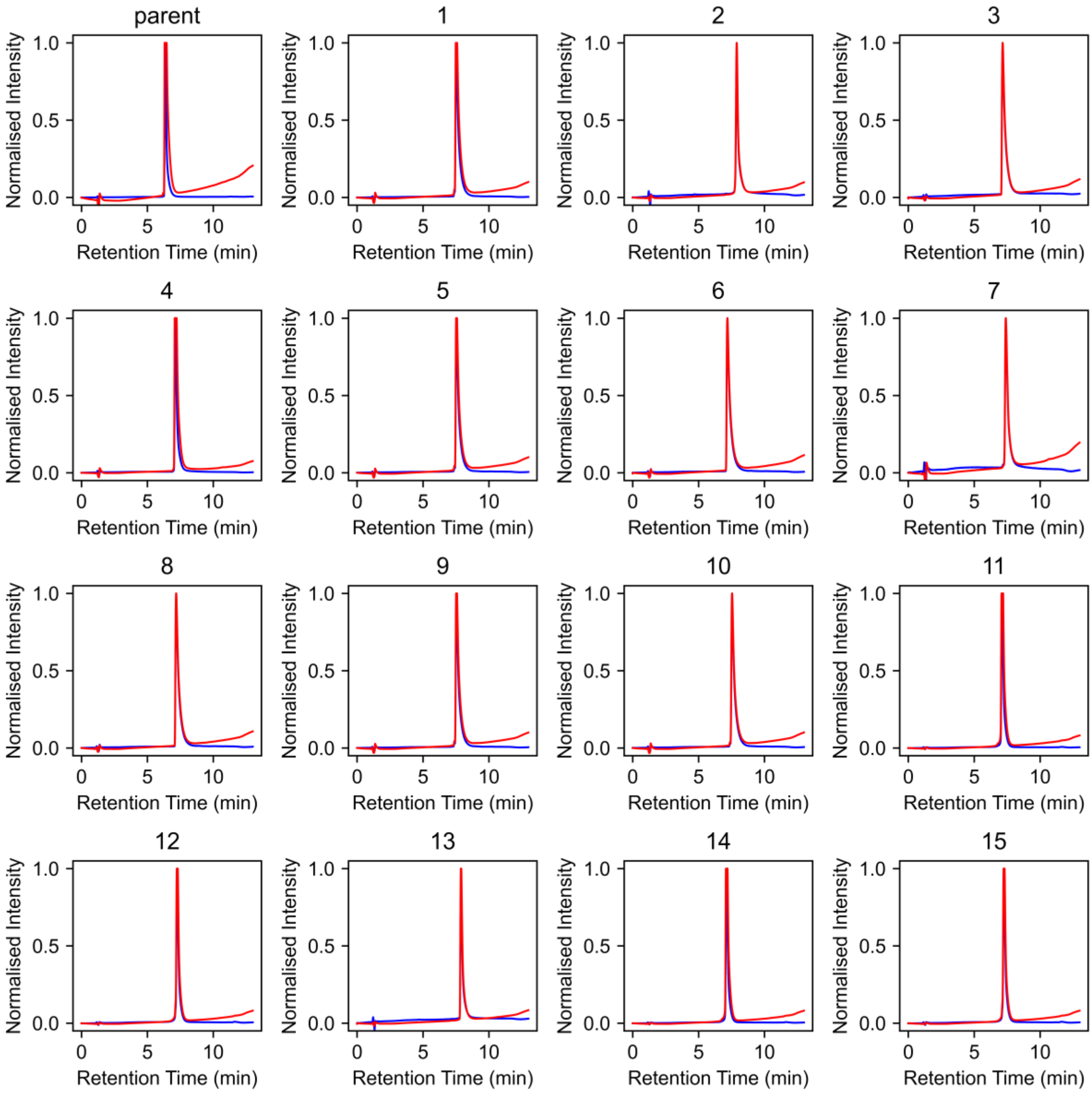


Figure S2 Analytical HPLC traces of *de novo* peptides described in this work.  Key: blue – 280 nm, red – 220 nm.

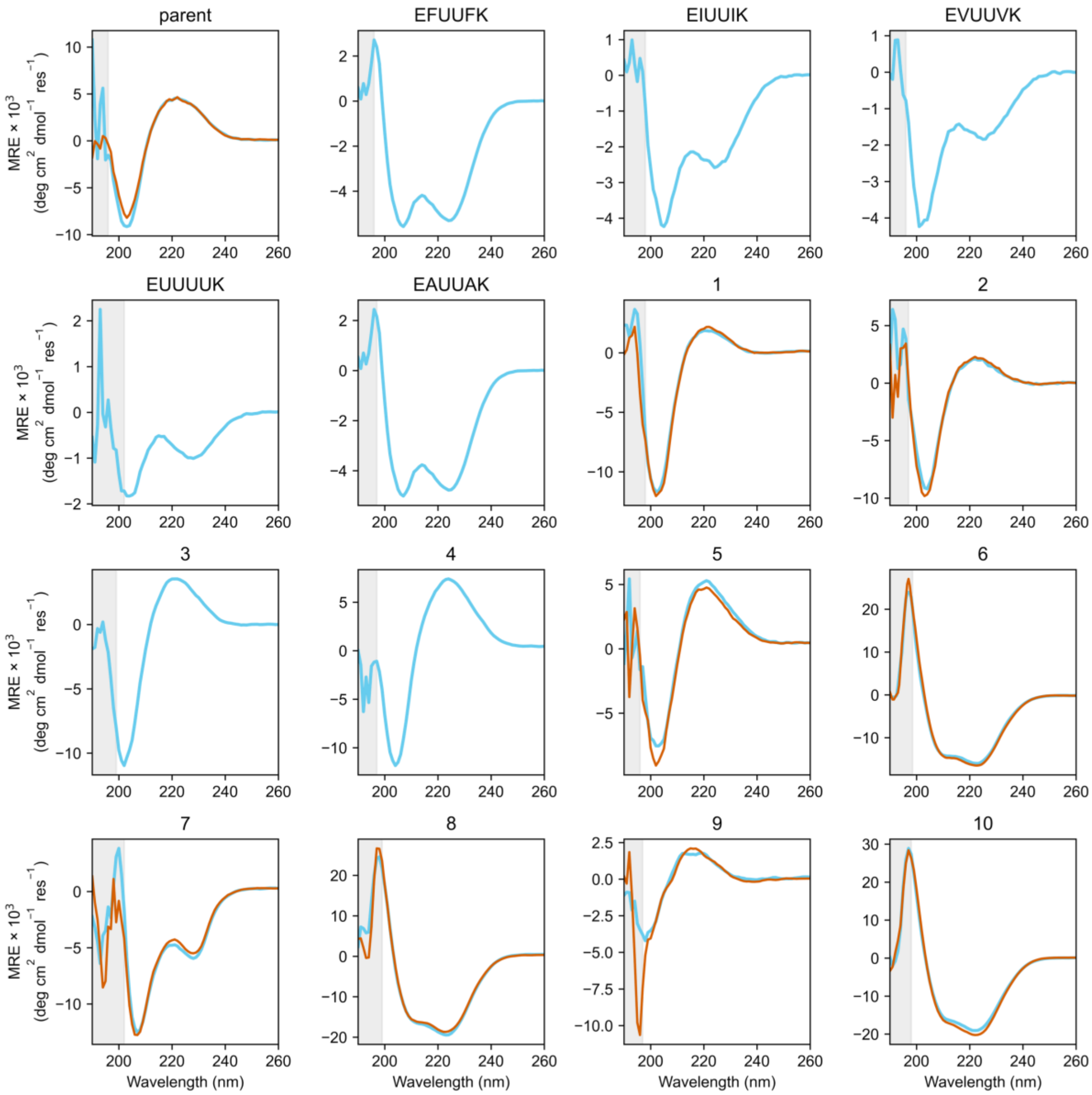

parent
EFUUFK
EIUUIK
EVUUVK
EUUUUK
EAUUAK
1
2
3
4
5
6
7
8
9
10
MRE × 10$^3$ (deg cm$^2$ dmol$^{-1}$ res$^{-1}$)
Wavelength (nm)


196

197

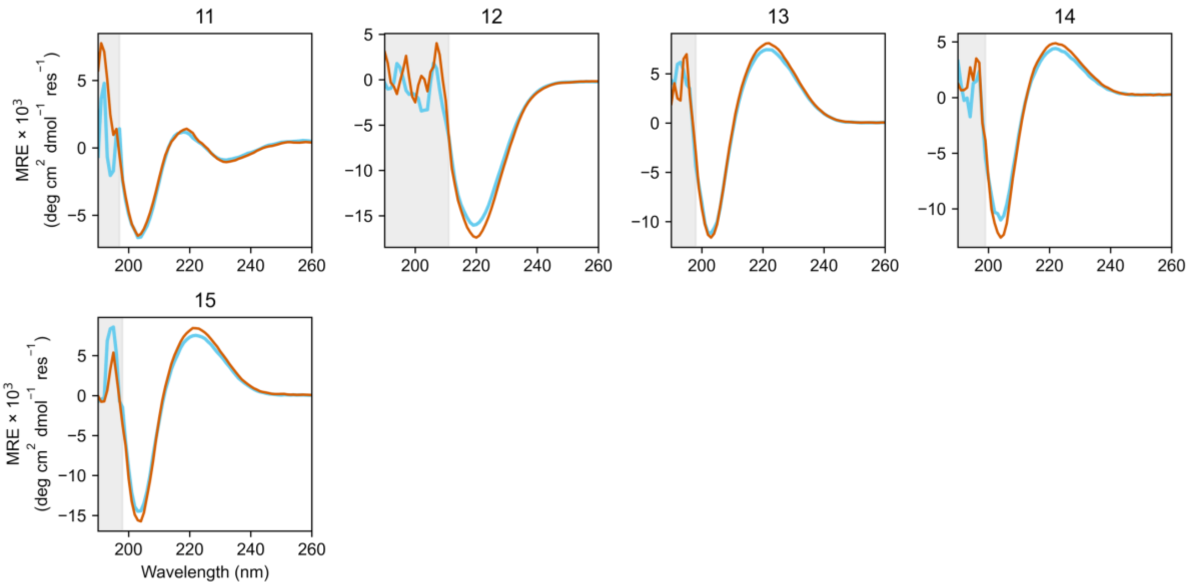


Figure S3 CD spectra of the *de novo* peptides discussed in this study. Summary of biophysical characterization can be found in Table S1. Conditions: 100 μM peptide assembly, PBS, pH 7.4. CD scans before the thermal ramps up (blue line) and after thermal ramps down (orange line).

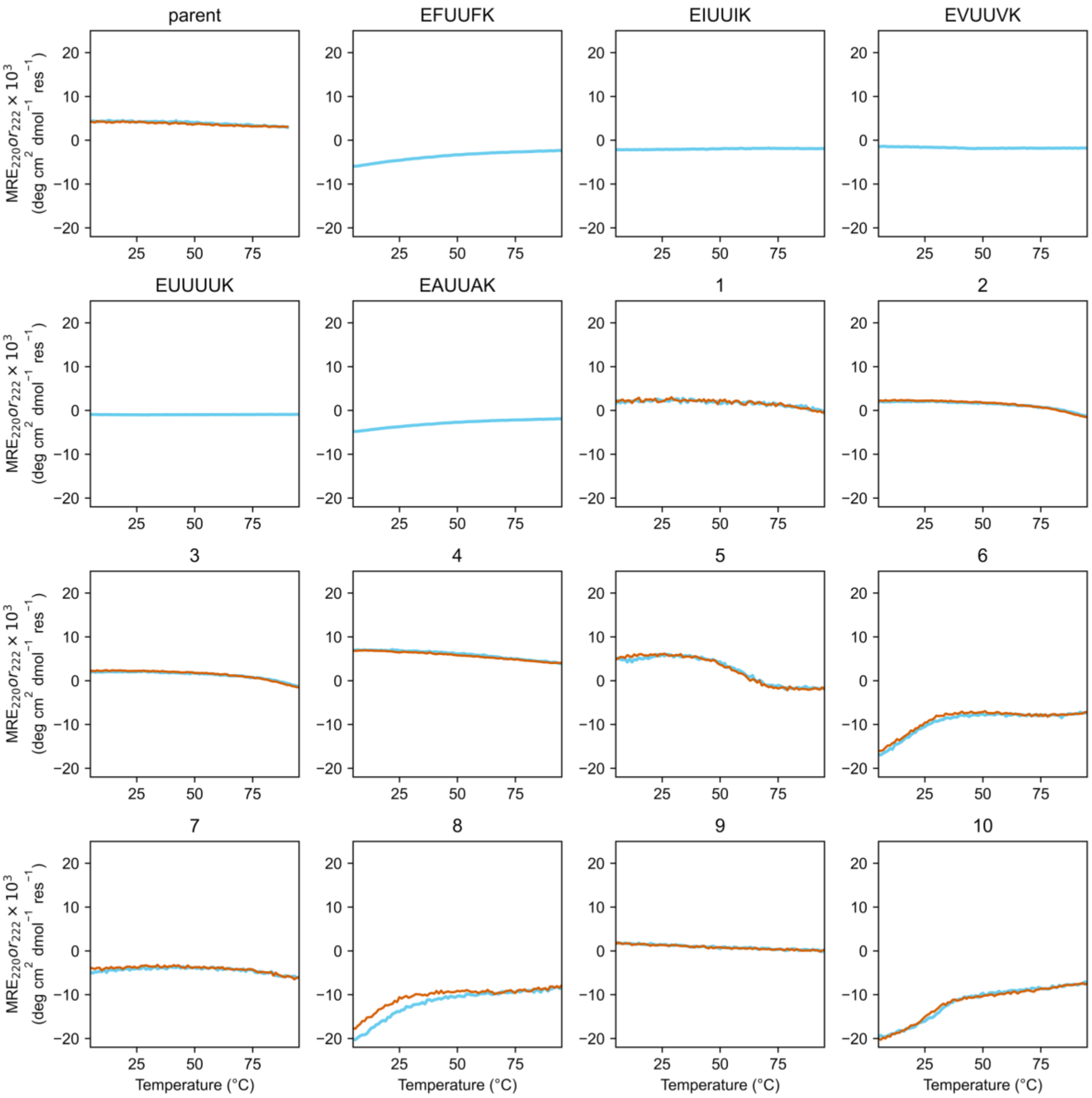
parent
EFUUFK
EIUUIK
EVUUVK
EUUUUK
EAUUAK
1
2
3
4
5
6
7
8
9
10
MRE220 or r222 × 10^3 (deg cm^2 dmol^−1 res^−1)
Temperature (°C)
20
10
0
−10
−20
25
50
75

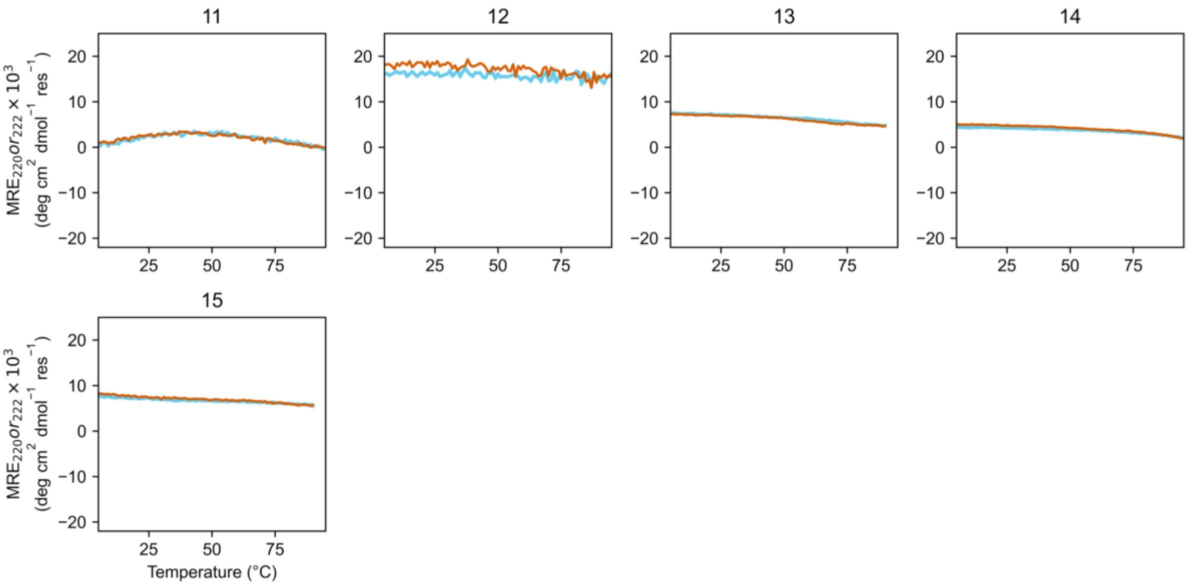


Figure S4 Variable-temperature CD spectra of the *de novo* peptides discussed in this study. Summary of biophysical characterization can be found in Table S1. Conditions: 100 µM peptide assembly, PBS, pH 7.4; 5 to 95 °C, 1 °C per minute ramp rate. Keys: thermal ramps up (blue line) and thermal ramps down (orange line).

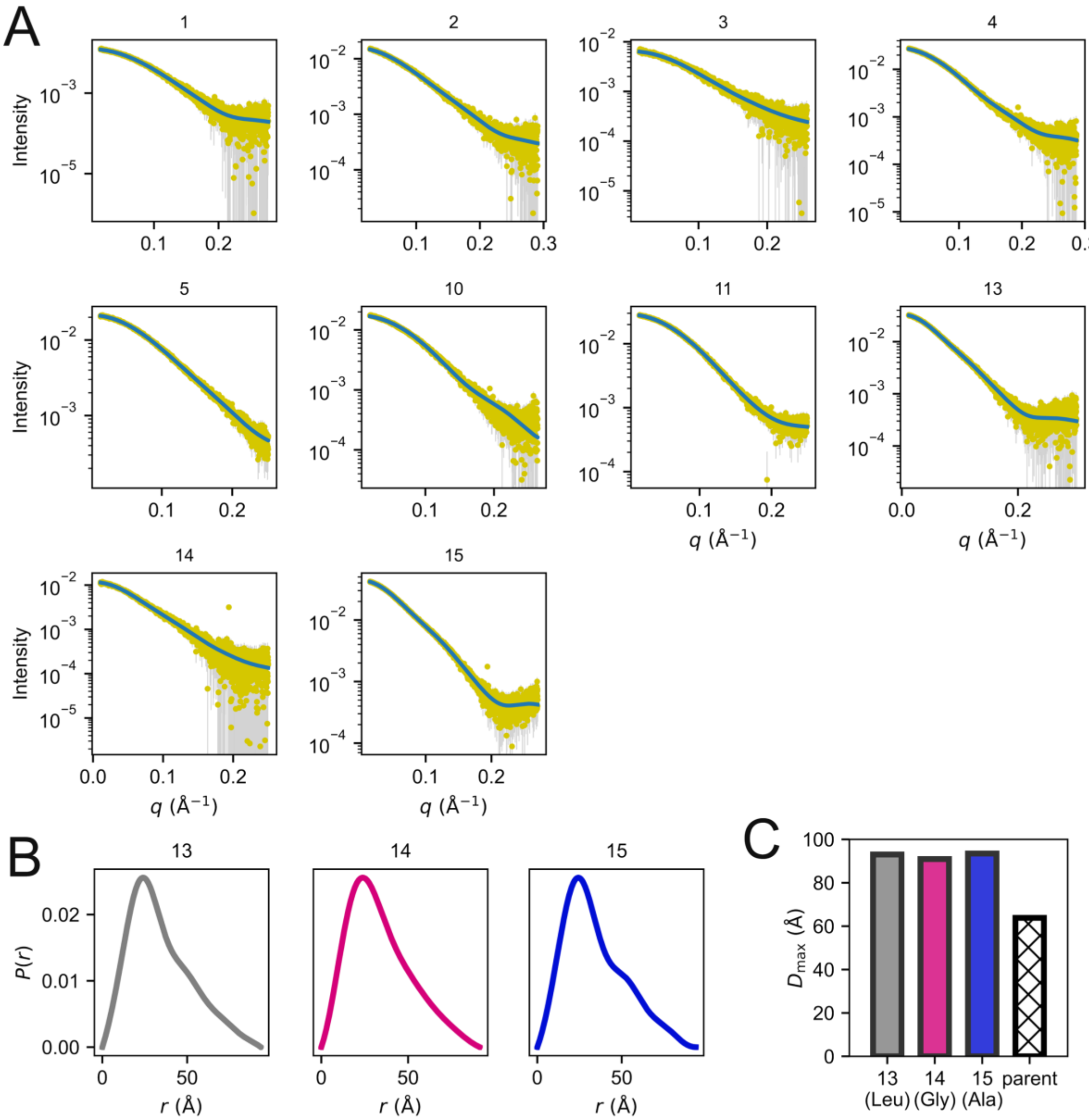


Figure S5 Small-angle X-ray scattering (SAXS) characterization of the *de novo* peptides discussed in this study. Summary of SAXS can be found in Table S2. (A) Experimental SAXS profiles (intensity vs. momentum transfer, q). Yellow points with grey error bars represent experimental scattering data; blue lines represent the corresponding fit generated in GNOM. Intensity is shown on a logarithmic scale. (B) Pair-distance distribution functions, P(r), calculated from the scattering profiles in (A) for constructs 13–15, highlighting differences in maximum particle dimension ($D_{max}$) compared to the parent resulting from the additional 3 hexads. (C) Comparison of $D_{max}$ values derived from the P(r) distributions in (B), alongside the parent construct (hatched bar) for reference. SAXS measurements were performed at 2.5 mg ml$^{-1}$ peptide in 20 mM Tris pH 8.0, 50 mM NaCl, 1% *v/v* glycerol.

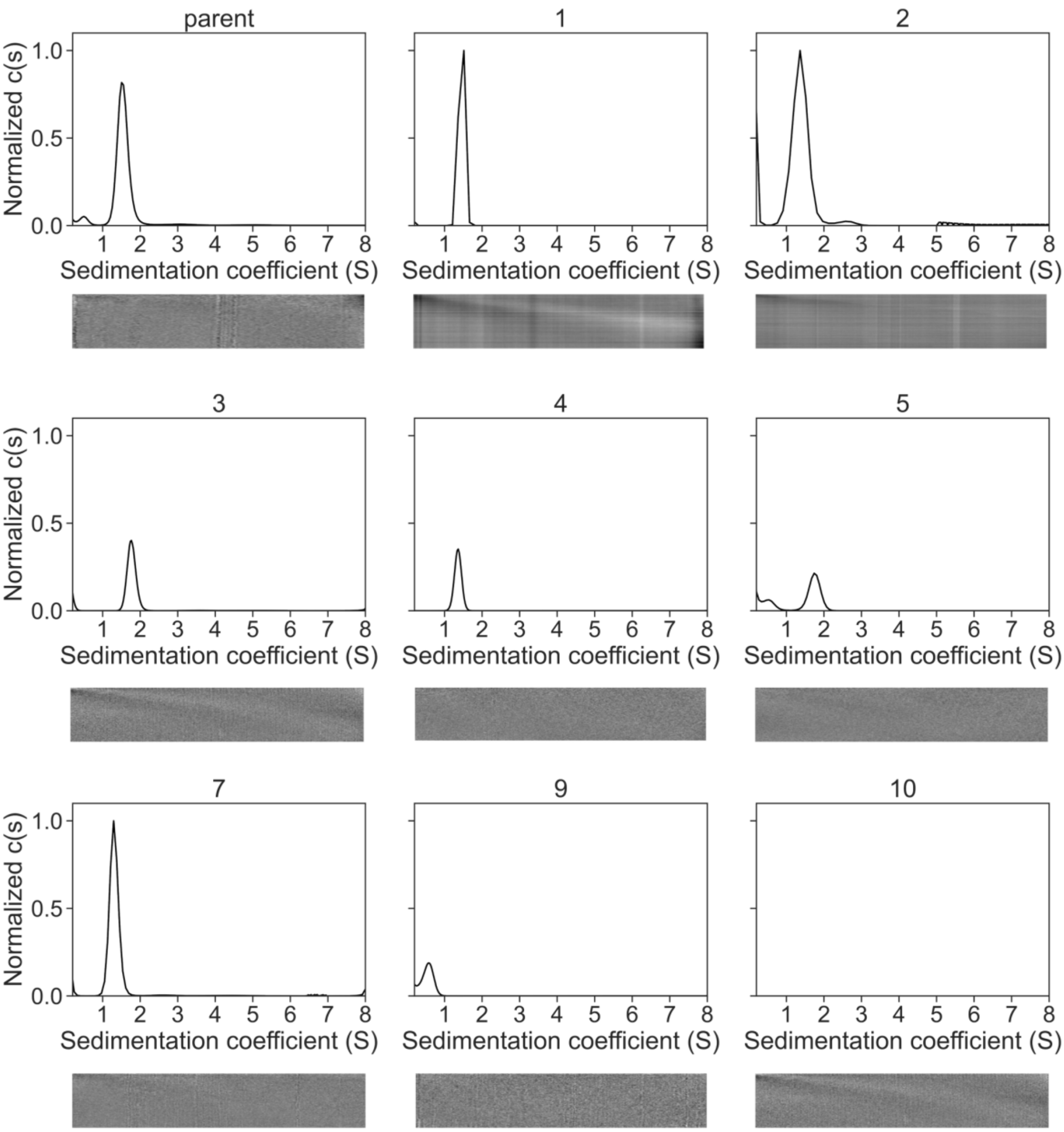
parent
1
2
3
4
5
7
9
10
Normalized c(s)
0.0
0.5
1.0
Sedimentation coefficient (S)
1 2 3 4 5 6 7 8

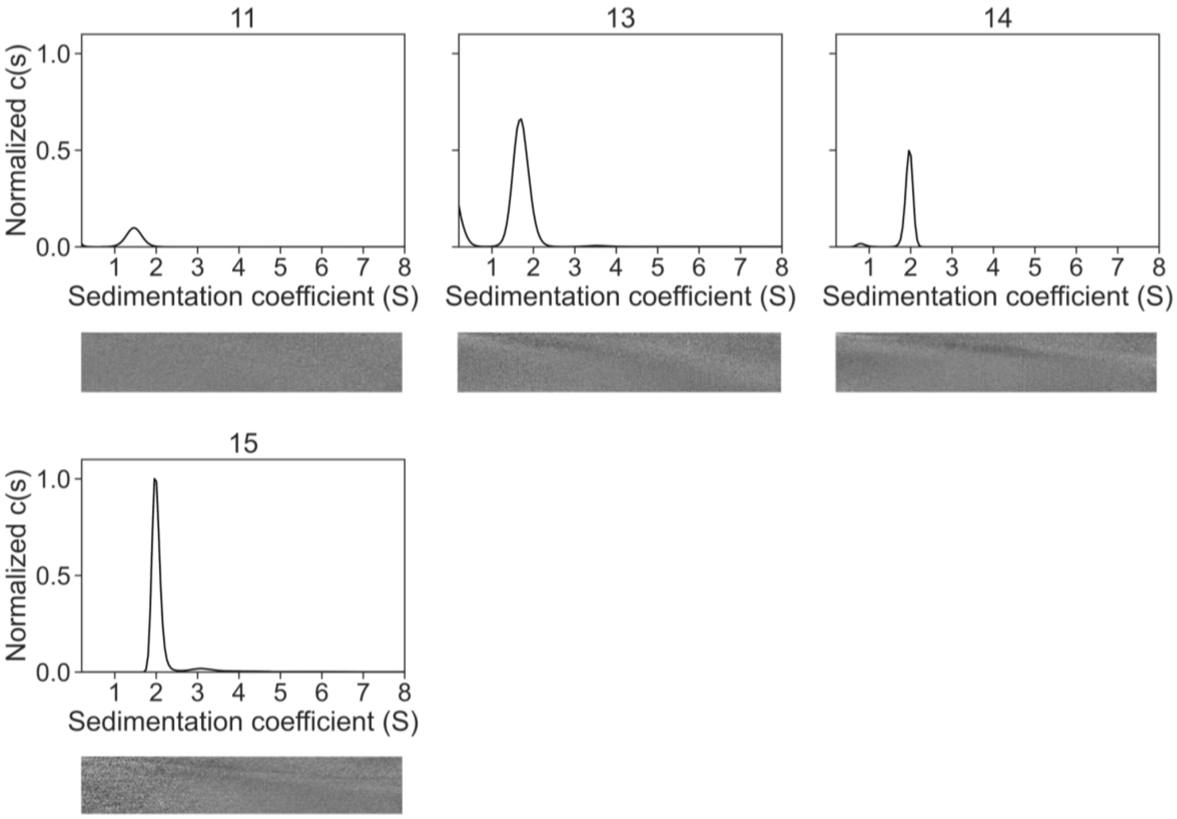


Figure S6 AUC-SV traces of the *de novo* peptides described in this work. Sequences for individual peptides can be found in Table 1. Summary of biophysical characterization can be found in Table S1. Residuals are shown as bitmaps below the fitted data. Conditions: 100 µM overall peptide assembly, PBS, pH 7.4, 20 °C, 50 krpm.

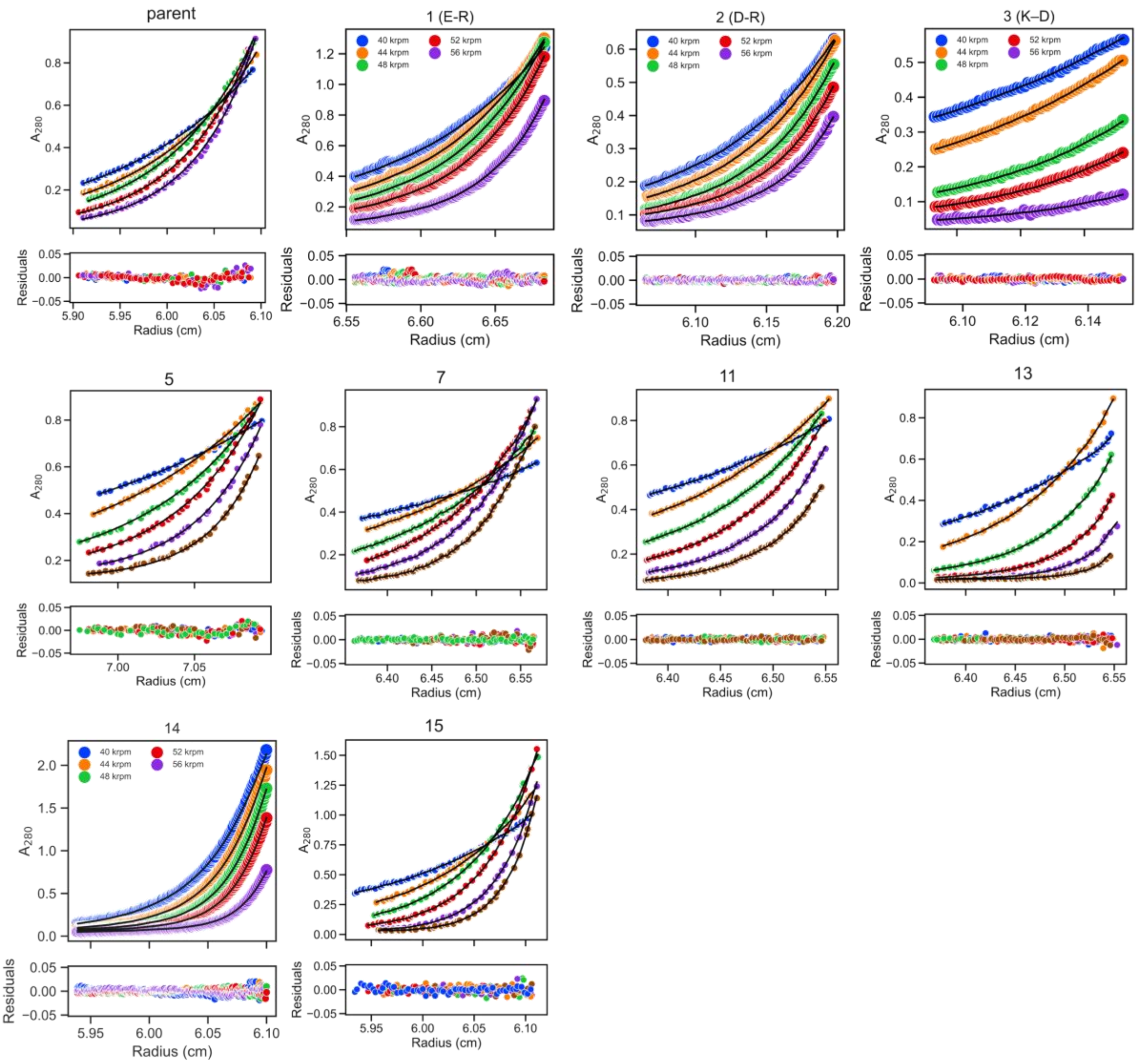


Figure S7 AUC-SE traces of the *de novo* peptides described in this work. Sequences for individual peptides can be found in Table 1. Summary of biophysical characterization can be found in Table S1. Conditions: 100 µM overall peptide assembly, PBS, pH 7.4, 20 °C, 20 – 45 krpm, or 40 – 56 krpm when specified. Key: 20 krpm, blue; 25 krpm, orange; 30 krpm, green; 35 krpm, red; 40 krpm, purple; 45 krpm, brown.

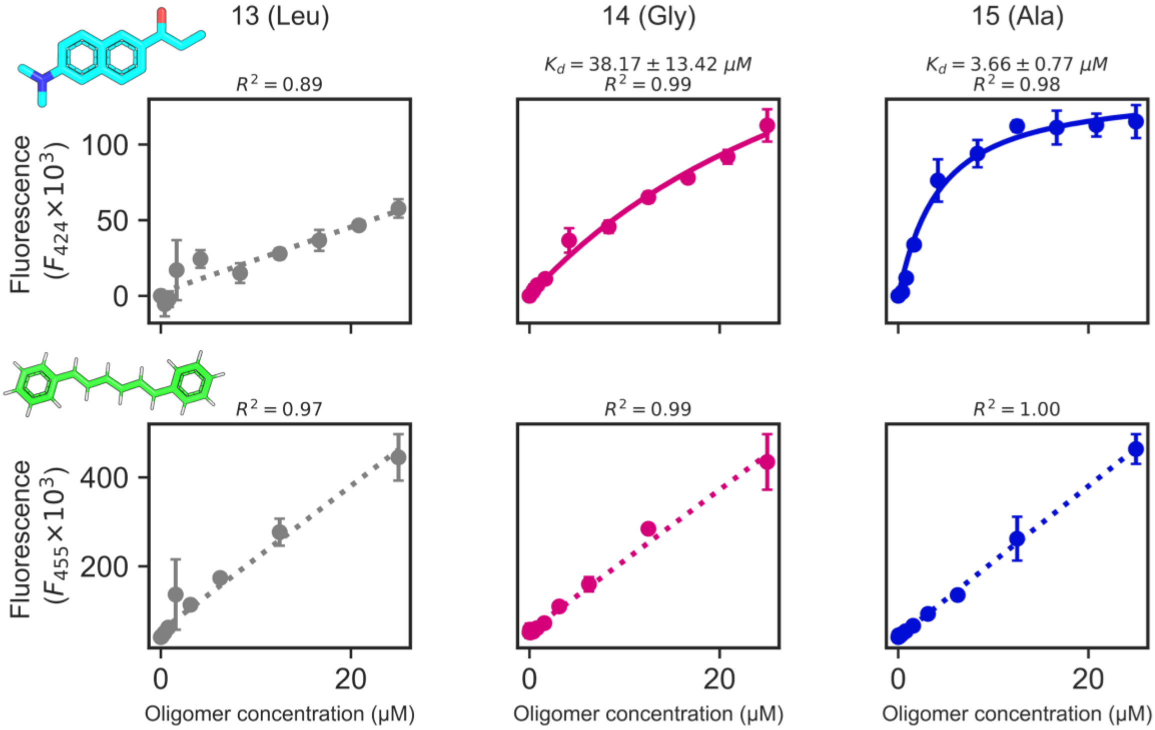


Figure S8 Fluorescence assay for prodan and DPH binding to peptides **13–15** described in this work. Prodan (top) and DPH (bottom) fluorescence measured as a function of oligomer concentration for the **13** (Leu), **14** (Gly), and **15** (Ala) variants. Data points represent mean fluorescence ± SD. Prodan binding data were fitted using a quadratic binding equation to account for ligand depletion, with the fitted $K_d$ and $R^2$ values indicated above each panel. Dotted fit lines indicate that data were fitted linearly. Conditions: 1 μM dye in PBS with 5% *v/v* DMSO; the concentration of peptide assembly (hexamer) was a serial dilution between 0 and 30 μM.

### References


(1) Zieleniewski, F.; Woolfson, D. N.; Clayden, J. Automated solid-phase concatenation of Aib residues to form long, water-soluble, helical peptides. *Chem. Commun.* **2020**, *56* (80), 12049–12052.

(2) Kuipers, B. J. H.; Gruppen, H. Prediction of Molar Extinction Coefficients of Proteins and Peptides Using UV Absorption of the Constituent Amino Acids at 214 nm To Enable Quantitative Reverse Phase High-Performance Liquid Chromatography−Mass Spectrometry Analysis. *J. Agric. Food. Chem.* **2007**, *55* (14), 5445–5451.

(3) Sreerama, N.; Woody, R. W. Computation and Analysis of Protein Circular Dichroism Spectra. In *Methods in Enzymology*, Vol. 383; Academic Press, 2004; pp 318–351.

(4) Lindner, W. A.; Brand, J. M. A coherent approach to the svedberg equation. *Biochem. Educ.* **1987**, *15* (2), 71–72.

(5) Dawson, W. M.; Martin, F. J. O.; Rhys, G. G.; Shelley, K. L.; Brady, R. L.; Woolfson, D. N. Coiled coils 9-to-5: rational de novodesign of α-helical barrels with tunable oligomeric states. *Chem. Sci.* **2021**, *12* (20), 6923–6928.

(6) Franke, D.; Gräwert, T.; Svergun, D. I. New features in ATSAS-4, a program suite for small-angle scattering data analysis. *J. Appl. Crystallogr.* **2025**, *58* (3), 1027–1033.

(7) Franke, D.; Petoukhov, M. V.; Konarev, P. V.; Panjkovich, A.; Tuukkanen, A.; Mertens, H. D. T.; Kikhney, A. G.; Hajizadeh, N. R.; Franklin, J. M.; Jeffries, C. M.; et al. ATSAS 2.8: a comprehensive data analysis suite for small-angle scattering from macromolecular solutions. *J. Appl. Crystallogr.* **2017**, *50* (4), 1212–1225.
